# The Growing Little Brain: Cerebellar Functional Development from Cradle to School

**DOI:** 10.1101/2024.10.12.617938

**Authors:** Wenjiao Lyu, Kim-Han Thung, Khoi Minh Huynh, Li Wang, Weili Lin, Sahar Ahmad, Pew-Thian Yap

**Affiliations:** Department of Radiology, University of North Carolina, Chapel Hill, NC, USA; Biomedical Research Imaging Center, University of North Carolina, Chapel Hill, NC, USA

## Abstract

Despite the cerebellum’s crucial role in brain functions, its early development, particularly in relation to the cerebrum, remains poorly understood. Here, we examine cerebellocortical connectivity using over 1,000 high-quality resting-state functional MRI scans of children from birth to 60 months. By mapping cerebellar topography with fine temporal detail for the first time, we show the hierarchical organization of cerebellocortical functional connectivity from infancy. We observe dynamic shifts in cerebellar network gradients, which become more focal with age while generally maintaining stable anchor points similar to adults, highlighting the cerebellum’s evolving yet stable role in functional integration during early development. Our findings provide the first evidence of cerebellar connections to higher-order networks at birth, which generally strengthen with age, emphasizing the cerebellum’s early role in cognitive processing beyond sensory and motor functions. Our study provides insights into early cerebellocortical interactions, reveals functional asymmetry and sex-specific patterns in cerebellar development, and lays the groundwork for future research on cerebellum-related disorders in children.

---

The cerebellum’s contributions to human language, emotional regulation, attention control, cognition, and working memory, beyond its widely acknowledged role in motor functions, have garnered increasing attention in recent years^1–3^. These diverse functions are believed to result from the cerebellum’s interaction with the cerebrum, positioning the cerebellocortical system as one of the most crucial circuits in the human brain^4^. Evolutionary anthropologists suggest that brain expansion in primates, including humans, is driven by the selective modular expansion of the cerebellocortical system^5^. Notably, compared to other primates, key regions associated with cognition in the human cerebellocortical system expand at the highest evolutionary rates^6^, indicating that this system may play a pivotal role in the evolution of human brain functions and may offer critical clues on the formation of higher-order cognitive functions in humans. Numerous studies, including those involving non-human primate experiments and human neuroimaging research, have confirmed the connections between the cerebellum and cortex^7^. Additionally, clinical studies^8, 9^ have shown that damage to the connections between the cerebellum and cortex during development may lead not only to cerebellar abnormalities but also impairment of cerebral cortical development, underscoring the importance of cerebellocortical connectivity in early neurodevelopment. However, there is still limited knowledge about typical cerebellocortical connectivity and how it develops in early human life.

Children progress through several distinct stages in early childhood—neonatal, infancy, toddlerhood, and preschool—each marked by rapid motor and cognitive development. In the neonatal stage, children develop basic reflexes and motor responses, such as grasping, head movements, and recognizing their mother’s voice. As they transition into infancy, they begin to acquire more complex motor skills, like rolling, sitting, and standing, while gradually displaying higher-order functions like recognizing familiar faces and objects, using simple gestures, and beginning to speak. The toddler stage sees children mastering walking, running, jumping, and throwing, which require coordination, balance, and motor planning. Additionally, they also experience a wider range of emotions, imitate behaviors, and increasingly use language for communication^10–15^. During the preschool years, children further refine these skills and begin to develop fine motor abilities, like drawing, as well as higher-order cognitive functions such as counting, categorization, language development, and storytelling^15, 16^. The rapid development of these higher-order functions, particularly before school age, distinguishes humans from other mammals. This intense developmental period has prompted researchers to study how the cerebellum’s unique connection to the cerebral cortex supports these advanced abilities.

The development of these motor and cognitive skills is supported by the maturation of neural substrates in the brain after birth, involving processes such as synaptogenesis (formation of new synapses), myelination (insulation of nerve fibers), programmed cell death (apoptosis), and synaptic pruning (removal of unnecessary synapses). These processes occur at different rates across various brain regions during early childhood^17, 18^. For example, the visual cortex, which processes sensory input, matures earlier than the prefrontal cortex, which is associated with decision-making and higher cognitive functions^17^. This difference in developmental timing shows that sensory and association regions—key components of the brain’s hierarchical organization—do not mature simultaneously. As a result, the organization and strength of cerebellocortical connections—the pathways linking the cerebellum and the cerebral cortex—are likely to differ significantly between children and adults. Understanding these differences can provide insights into how the brain supports the rapid development of complex sensorimotor and cognitive abilities during early childhood.

Over the past decade, functional magnetic resonance imaging (fMRI) has been used to study the functional connectivity between the cerebellum and cortical networks. Advances in this field have revealed the macroscale functional organization of the adult cerebellum based on its connections with cortical regions. Key findings include but are not limited to the cerebellum’s functional division into primary sensorimotor and supramodal zones^19^; its contralateral connectivity with the cerebral cortex^20^; the presence of multiple topographically organized somatomotor representations^20^; the involvement of cerebellar subregions in both integrative and segregative functions^21^; the contribution of phylogenetically recent regions, Crus I and Crus II, to connections with higher-order cognitive networks^22^; and a functional hierarchy along the central axis of motor and non-motor organization in the cerebellum^23–25^. Additionally, detailed functional mapping of the cerebellum has been achieved at both the group^20^ and individual^26^ levels.

Despite these advances in understanding cerebellocortical functional connectivity in adults, research on these connections in young children is still limited^27, 28^. This gap in knowledge is particularly concerning given that altered cerebellocortical functional connectivity has been observed in early-onset neurodevelopmental disorders such as autism spectrum disorder (ASD) and attention deficit hyperactivity disorder (ADHD)^29–34^. Therefore, elucidating typical cerebellocortical functional connectivity during early development is essential for establishing a foundational understanding that can inform the interpretation of these abnormalities and their broader implications on neurodevelopmental disorders. Several fundamental questions about early cerebellar development remain unanswered:

- When does the cerebellum first begin to contribute to higher-order functions?
- How do the patterns of cerebellar primary motor function evolve during early development?
- At what age do children’s cerebellar functions resemble those of adults?
- How does cerebellum development differ between male and female children?
- Is there a difference in cerebellar development between the left and right hemispheres, and if so, when does this lateralization commence?
- What is the spatiotemporal pattern of early cerebellar development?
- How does the functional hierarchy of the cerebellum evolve throughout early development?

To answer these questions, we mapped the functional connectivity patterns between the cerebellum and cerebral resting-state networks (RSNs) in children from birth to 60 months, using over 1,000 fMRI scans from 275 participants in the Baby Connectome Project (BCP)^35^. We examined the development and organization of cerebellocortical functional connectivity over time, focusing on the establishment and timing of higher-order network connections. Additionally, we explored spatiotemporal patterns, lateralization, and sex differences in cerebellocortical functional connectivity, contributing to a broader understanding of this critical period of early development.

## RESULTS

We applied group independent component analysis (GICA) to preprocessed fMRI data to generate 40 RSNs, from which we selected 30 cortical RSNs to ensure full coverage of the cerebral cortex. Using these RSNs, we evaluated cerebellocortical functional connectivity through partial correlation analysis with the cerebellum. These 30 RSNs were categorized into eight large-scale networks: Sensorimotor Network (SMN), Auditory Network (AUD), Visual Network (VIS), Salience Network (SN), Ventral Attention Network (VAN), Default Mode Network (DMN), Executive Control Network (ECN), and Dorsal Attention Network (DAN) (Figure 1). Each RSN was named based on its affiliation with a large-scale network and its predominant location in the brain (Table S1). To clarify the principles of cerebellocortical organization, we grouped the SMN, AUD, and VIS as primary networks, and the SN, VAN, DMN, ECN, and DAN as higher-order networks, depending on whether they are primarily engaged in sensory and motor functions or higher-order cognitive functions.

**Fig. 1.**
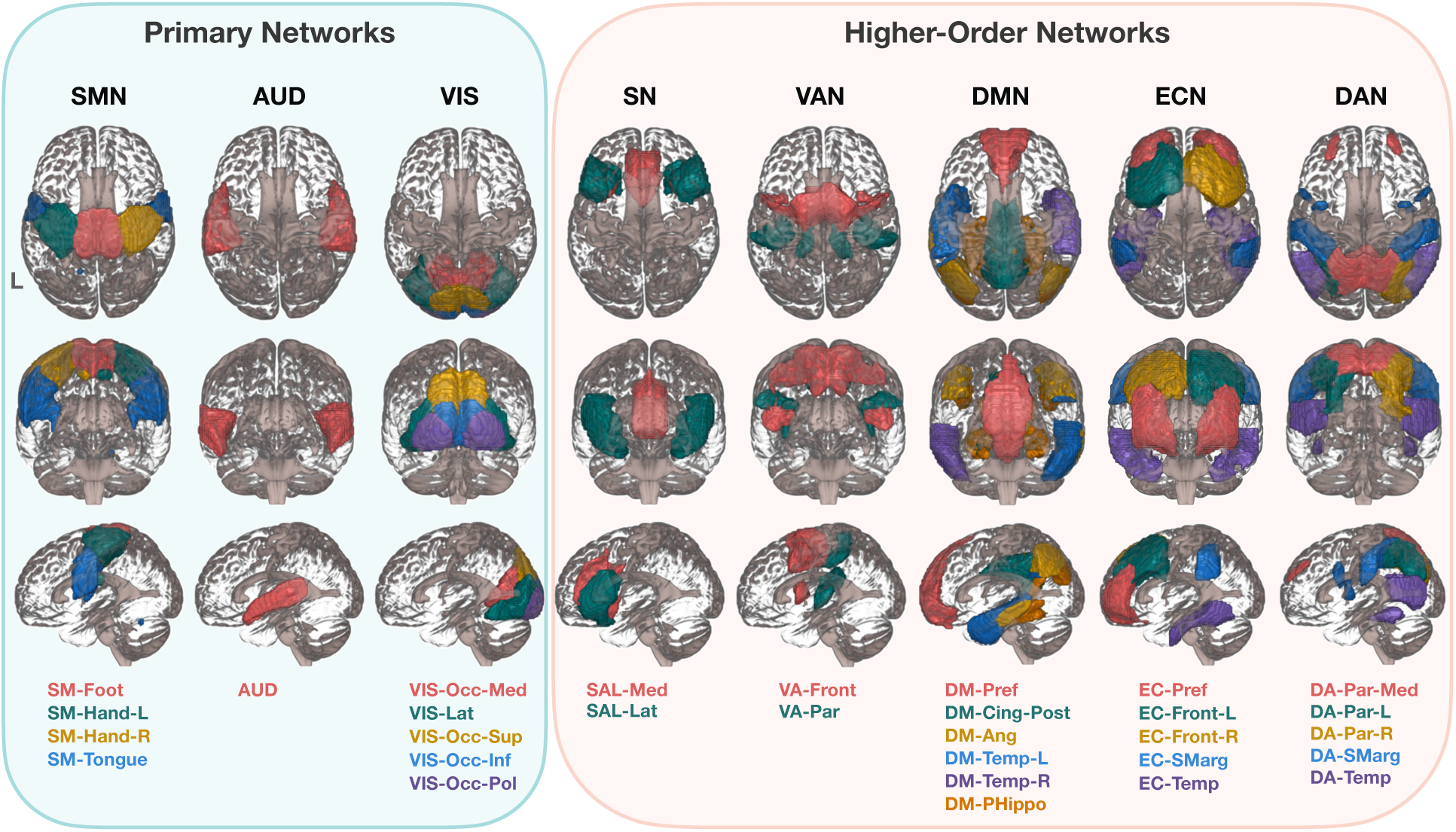
Cortical networks. Except the VIS and the DAN, which are presented from superior to inferior, posterior to anterior, and left to right, all networks are displayed from superior to inferior, anterior to posterior, and left to right. SMN: sensorimotor network; AUD: auditory network; VIS: visual network; SN: salience network; VAN: ventral attention network; DMN: default mode network; ECN: executive control network; DAN: dorsal attention network; SM: sensorimotor; L: left; R: right; Occ: occipital; Med: medial; Lat: lateral; Sup: superior; Inf: inferior; Pol: pole; SAL: salience; VA: ventral attention; DM: default mode; Pref: prefrontal; Cing: cingulate gyrus; Post: posterior; Ang: angular gyrus; Temp: temporal; PHippo: parahippocampus; EC: executive control; Front: frontal; SMarg: supramarginal gyrus; DA: dorsal attention; Par: parietal.

### Cerebellocortical Functional Connectivity

We first verified the veracity of our framework in capturing cerebellar functional organization by localizing the sensorimotor representations of the foot, hand, and tongue^20, 26^. We evaluated the functional connectivity between the SM-Foot, SM-Hand, SM-Tongue and the cerebellum and observed that sensorimotor representations in the cerebellum during early childhood exhibit organizational characteristics similar to those in adults. Specifically, in the anterior lobe, the representation is inverted, with the foot anterior to the hand and tongue, whereas in the posterior lobe, the representation is upright, with the tongue anterior to the hand and feet (Figure 2a). Using the spatially unbiased atlas template (SUIT) software package developed by Diedrichsen and colleagues^36^, we confirmed that the SMN representations are situated at expected positions on the flat map (Figure 2b), gauging based on lobular dermacation (Figure 2c). Subsequently, we depicted the spatial and temporal changes in cerebellocortical functional connectivity using network-specific flat maps (Figure 3 and Figure S2) and age-related trajectories (Figure S3).

**Fig. 2.**
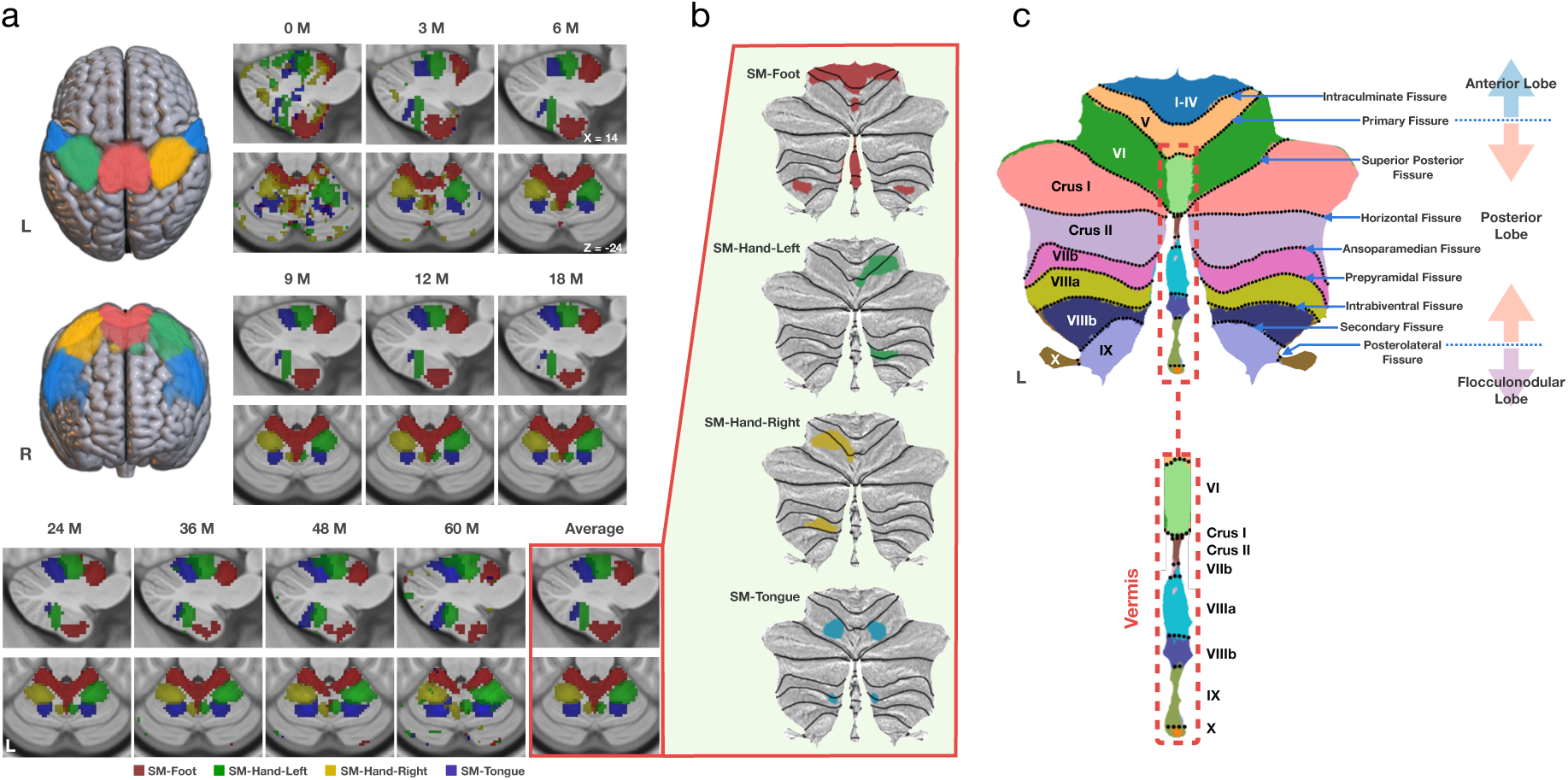
Representations of sensorimotor networks. **a**, Sensorimotor networks and their representations in the cerebellum across age, along with the average sensorimotor representation in MNI space, thresholded at *z* = 3.5. **b**, Flat maps of the average sensorimotor representations in the cerebellum, thresholded at *z* = 3.5. **c**, Lobular, fissural, and lobar annotations on the cerebellar flat map^36, 37^.

**Fig. 3.**
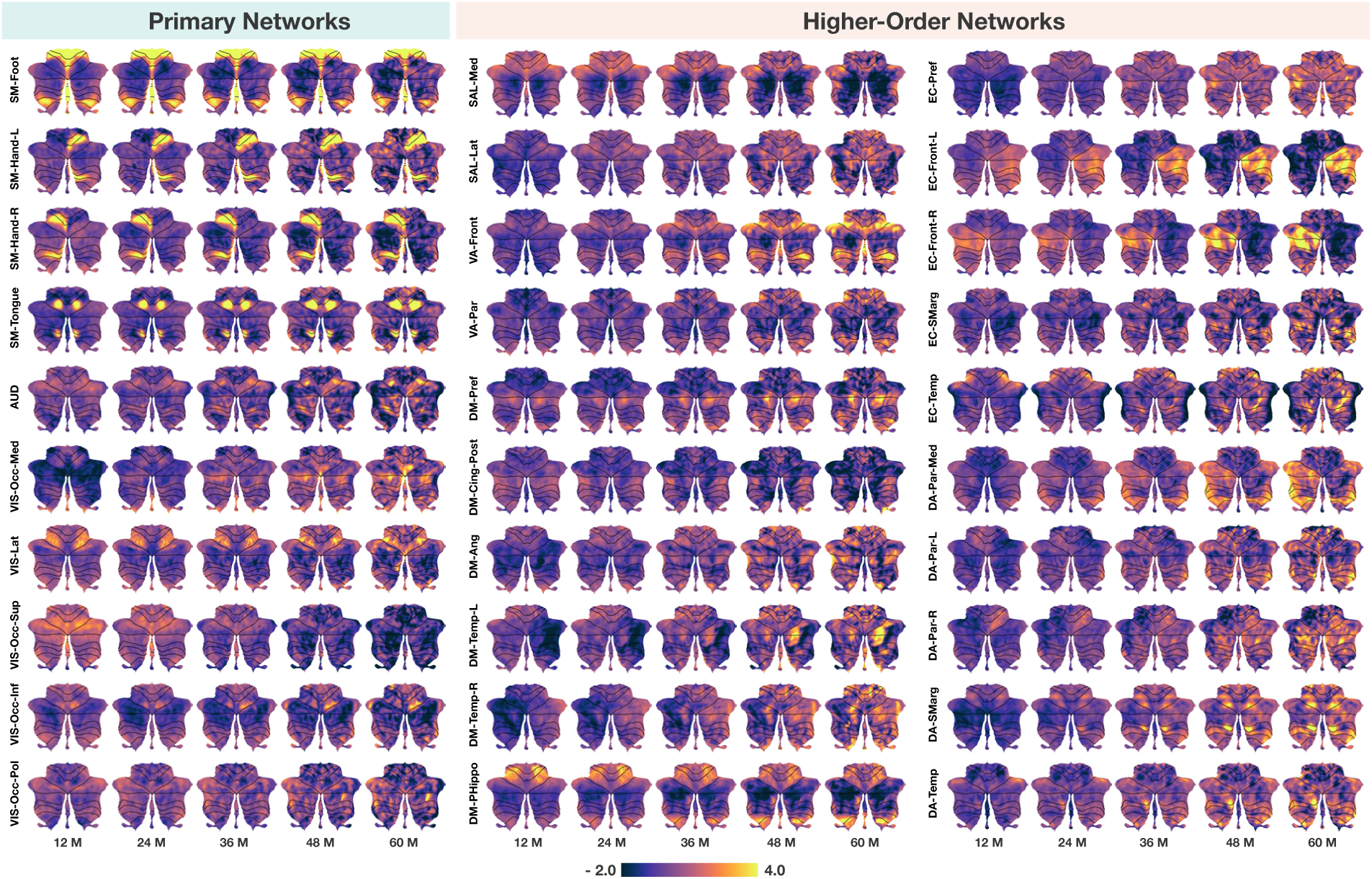
Cerebellar functional flat maps. Spatiotemporal patterns of connectivity (z-transformed) between the cerebellum and primary networks and higher-order networks at 12 months, 24 months, 36 months, 48 months, and 60 months. Values outside the range of −2.0 to 4.0 are capped for clarity.

For primary networks (Figure 3, Primary Networks), heightened connectivity was observed between several cerebellar regions, encompassing Lobules I-VI and VIII, and the SMN in early childhood, in line with previous studies on adults^20, 26, 38^. Specifically, Lobules I-V, the lateral aspect (away from the vermis) of Lobule VIII, and Vermis VI-VIII exhibit robust connectivity with SM-Foot. Lobules V-VI and the medial aspect (towards the vermis) of Lobule VIII exhibit strong connectivity with SM-Hand. As expected, the right cerebellum exhibits more pronounced connectivity with left SM-Hand, and vice versa. Additionally, Lobule VI and the medial aspect of Lobule VIIIa exhibit strong connectivity with SM-Tongue. Interestingly, unlike previous studies focused on adults, connectivity between the cerebellum and AUD is evident in early childhood, particularly during the first few months of life (Figure S2, Primary Networks). Additionally, we identified robust connectivity between the VIS and multiple cerebellar regions, including Lobule VI, Crus I, and the vermis, encompassing a broader spatial extent than previously reported in the literature^20, 26, 39^. Notably, cerebellar regions exhibiting strong connectivity to VIS shrink with age, mainly localizing at the inner aspect of Lobule VI and Vermis VI-VII from 48 months (Figure S2, Primary Networks), potentially indicating network specialization refinement.

Consistent with prior research on adults, cerebellar regions exhibiting heightened connectivity between higher-order networks are predominantly located in Lobules VI, Crus I, Crus II, IX, and X (Figure 3 and Figure S2, Higher-Order Networks). Despite the cerebellum’s typically weak and diffuse connectivity with most higher-order networks, strong connections were observed between the cerebellum and certain higher-order RSNs such as DM-PHippo, DM-Pref, bilateral EC-Front, EC-Smarg and EC-Temp, which were evident even at birth. Additionally, a trend of predominant contralateral connectivity was observed between the cerebellum and ECN-Front. Specifically, the right cerebellum is predominantly connected with left ECN-Front, whereas the left cerebellum is predominantly connected with right ECN-Front. The cerebellum generally exhibits stronger connectivity with primary networks compared to higher-order networks, especially during the first year of life, as demonstrated by the trajectories of the peak connectivity (Figure S3a). Biweekly growth rates indicate that connectivity between the cerebellum and the SM-Hand-L (primarily associated with right-hand motor functions) and SM-Tongue (primarily associated with tongue motor functions) increases with age, while connectivity with most primary networks remains stable or declines. Conversely, connectivity between the cerebellum and most higher-order networks tends to increase with age (Figure S3b). Together, these findings reveal dynamic changes in the spatial distribution and strength of cerebellar connectivity with RSNs during early childhood.

### Cerebellar Functional Topography

To better capture the spatiotemporal evolution of cerebellocortical functional connectivity across early childhood, we employed a winner-take-all approach to identify the network with the strongest connection to each cerebellar voxel^20, 27, 43, 44^, generating parcellation maps at three levels of granularity: (i) coarse granularity with two networks (primary and higher-order networks), (ii) medium granularity with eight networks (large-scale networks), and (iii) fine granularity with thirty networks (resting-state networks) (Figure 4a).

**Fig. 4.**
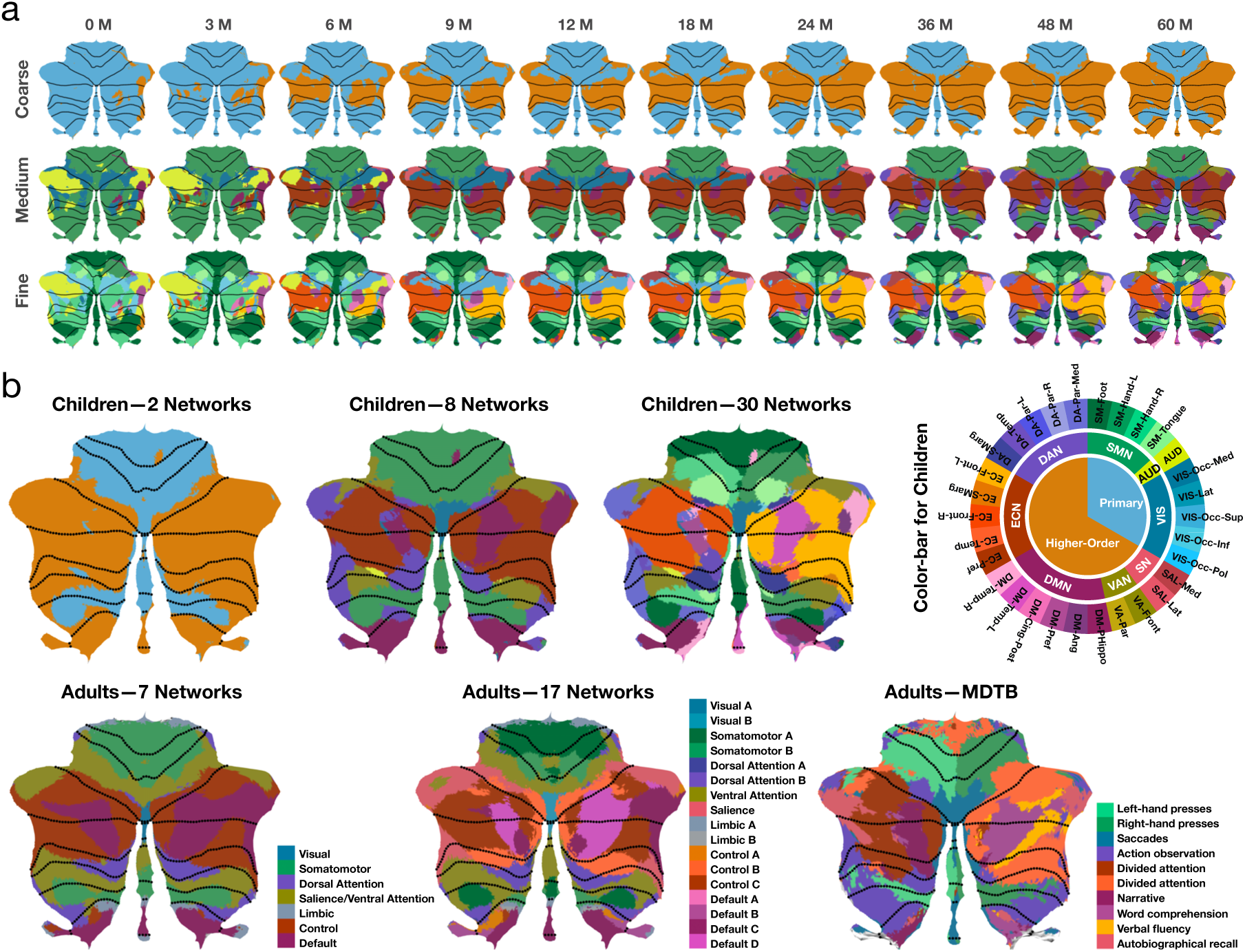
Comparison of parcellation maps between children and adults. **a**, Spatiotemporal patterns of cerebellar functional topography in early children at coarse, medium and fine granularity. **b**, Parcellation maps for children at 60 months, Buckner et al.’s 7 and 17 cerebellar resting-state networks^40^, and the MDTB^41^ parcellation map for adults.

At coarse granularity (Figure 4a, first row), we observed that although the cerebellum is predominantly connected to primary networks during the first few months after birth, regions in the posterior cerebellar lobe—particularly Crus I, Crus II, Lobule VIIb, and Lobule VIII—as well as those in the flocculonodular lobe that connect to primary networks shrink over time. By 60 months, cerebellar regions linked to primary networks are primarily restricted to Lobules I-VI, Lobule VIII, and Vermis VI-VIII, exhibiting an adult-like pattern. Regions in the posterior cerebellar lobe, particularly Crus I and Lobule VIII, connected to primary networks shrink with age. In contrast, regions in the posterior and flocculonodular lobes assigned to higher-order networks expand with age. Given that primary networks align with cortical sensory regions, while higher-order networks align with association regions, these findings suggest a developmental hierarchy stratified by the sensory-association (S-A) axis^45^.

At medium granularity (Figure 4a, middle row), regions in the anterior lobe connected to the SMN remain relatively stable throughout early childhood, while regions in the posterior and flocculonodular lobes connected to the SMN gradually shrink and concentrate in Lobule VIII by 60 months. Despite the presence of large cerebellar regions connected to the AUD in the first few months after birth, these regions notably decrease from 6 months onward, disappearing almost entirely before reappearing in small patches around 36 months. Initially, most cerebellar regions connected to the VIS are located in Lobule VI and Crus I. However, as age increases, the areas of Lobule VI and Crus I connected to VIS decrease, and by 60 months, the cerebellar regions connected to VIS are mainly concentrated at Vermis VI. Notably, Crus I, Crus II, and Lobule VIIb of the cerebellum are predominantly connected to the ECN, DMN, DAN, VAN, and SN, networks associated with higher-order functions, for most of early childhood (starting around 6 months).

Functional parcellation at fine granularity (Figure 4a, bottom row) provides a more detailed depiction of the topography of each RSN in the cerebellum, capturing subtle spatiotemporal changes that might be overlooked at medium granularity. For example, the fine-granularity parcellation revealed characteristic contralateral cerebellar connectivity with cortical networks, which extends beyond the SMN-related RSNs. Specifically, the right cerebellum tends to connect to SM-Hand-L, DM-Temp-L, EC-Front-L, and DA-Par-L, whereas the left cerebellum tends to connect to SM-Hand-R, DM-Temp-R, and EC-Front-R. Notably, RSNs within the same large-scale network are often topographically adjacent in the cerebellum, even if they are spatially distant in the cerebral cortex. For example, EC-Front, EC-Temp, and EC-SMarg, which belong to the same large-scale network, the ECN, are topographically non-adjacent in the cortex but are adjacent in the cerebellum at 60 months, highlighting the cerebellum’s pivotal role in integrating brain function^27, 46^.

We also compared the cerebellar functional topography maps obtained from 60-month-old children with Buckner et al.’s cerebellar resting-state networks^40^ and the multidomain task battery (MDTB) cerebellar parcellation maps of adults^41^ (Figure 4b). Note that the MDTB parcellation map, derived from a set of tasks^41^, is labeled according to each region’s most defining function, although the region may be associated with multiple functions. At 60 months, the cerebellar functional topography of primary and higher-order networks in children exhibits a distinct hierarchical organization (Figure 4b, Children – 2 Networks). Regions predominantly connected to primary networks are mainly situated within Lobules I-VI and the medial aspect of Lobule VIII, wheareas regions dominated by higher-order networks are primarily located in Lobule VII (including Crus I, Crus II, and Lobule VIIb), the lateral aspect of Lobule VIII, Lobule IX and Lobule X, closely resembling the functional organization principles of adults, particularly the “double motor and triple nonmotor representation”^23, 24^ or “three-fold organization” pattern^47^.

From 36 to 60 months, the cerebellar functional topography of the eight large-scale networks in children (Figure 4a, middle row) progressively mirrors that of the seven resting-state functional networks in adults (Figure 4b, Adults – 7 Networks) with the following pattern of cerebellocortical functional connectivity:

- SMN – Lobules I-VI, Lobule VIII, and Vermis VIII.
- AUD – Small regions in Lobules VI and VIIb.
- VIS – Lobule VI and Vermis VI.
- SN – Lobule VI and Crus I.
- VAN – Lobule VI Crus I and Lobule VIII.
- DMN – Crus I, Crus II, Lobules IX-X, Vermis IX, and Vermis X.
- ECN – Crus I, Crus II, and Lobule VIIb.
- DAN – Crus I, Lobule VIIb, Lobule VIII, Lobule IX, and Lobule X.

In 60-month-old children (Figure 4b, Children – 8 Networks), the cerebellar connectivity topography resembles that of adults but has not yet fully matured. Compared to adults, children show greater involvement of the SMN, AUD, and VIS. Notably, they also exhibit increased engagement with the DAN but reduced involvement with the VAN and DMN. These findings suggest a developmental shift toward greater functional specialization as the cerebellocortical system matures, aligning with the evolving cognitive and sensorimotor demands of early childhood.

It is perhaps unsurprising that the cerebellar functional parcellation map of 60-month-old children resembles the parcellation map of adults based on resting-state functional networks (Figure 4b, Adults – 7 Networks and 17 Networks). Interestingly, the cerebellar functional parcellation map of 60-month-old children at fine granularity (Figure 4b, Children – 30 Networks) is similar to the adult MDTB parcellation map (Figure 4b, Adults – MDTB). Specifically, the children’s SM-Hand-R region largely overlaps with the “Left-hand Presses” region in the adult MDTB map, the SM-Hand-L region overlaps with the “Right-hand Presses” region, the VIS-Occ-Med region overlaps with the “Saccades” region, and the DA-Par-Med region overlaps with the “Action Observation” region. Additionally, the DM-Pref region corresponds with the “Narrative” region, and the DM-Temp-L region corresponds with the “Word Comprehension” region in the adult MDTB map. These findings suggest that early cerebellar functional organization in children is closely linked to specialized tasks, indicating that the cerebellum may develop complex functional mappings earlier than previously thought.

Collectively, these findings demonstrate the dynamic evolution of cerebellar functional topography in early childhood, reflecting the increasing alignment with adult patterns as cognitive development and sensorimotor integration progress.

### Cerebellar Functional Gradients

Macroscale gradients of functional connectivity organize systematic information into abstract representations, providing valuable insights into how function varies across space^45, 48^. Using gradient-based analysis, Guell and colleagues have established the spatial macroscale gradients of the cerebellum in adults^23, 24, 49^. However, the emergence and development of functional gradients in the cerebellum during childhood have not been well-explored, despite the rapid neurodevelopment occurring during this critical period. To address this gap, we employed LittleBrain^42^ to generate cerebellar functional gradient maps, complementing the functional parcellation maps by capturing subtle and gradual spatial changes in cerebellar function. LittleBrain creates a two-dimensional map of cerebellar voxels, visualizing functional gradients with each axis representing a principal gradient: Gradient 1, which transitions from primary (motor) to transmodal (DMN, task-unfocused) regions, and Gradient 2, which shifts from task-unfocused (mind-wandering, goal-undirected thought) to task-focused (attentive, goal-directed thought) processing areas. Cerebellar voxels that are adjacent in the gradient map share similar functional connectivity patterns. The gradient map derived from the adult cerebellar functional parcellation^40, 42^ served as the reference (Figure 5a).

**Fig. 5.**
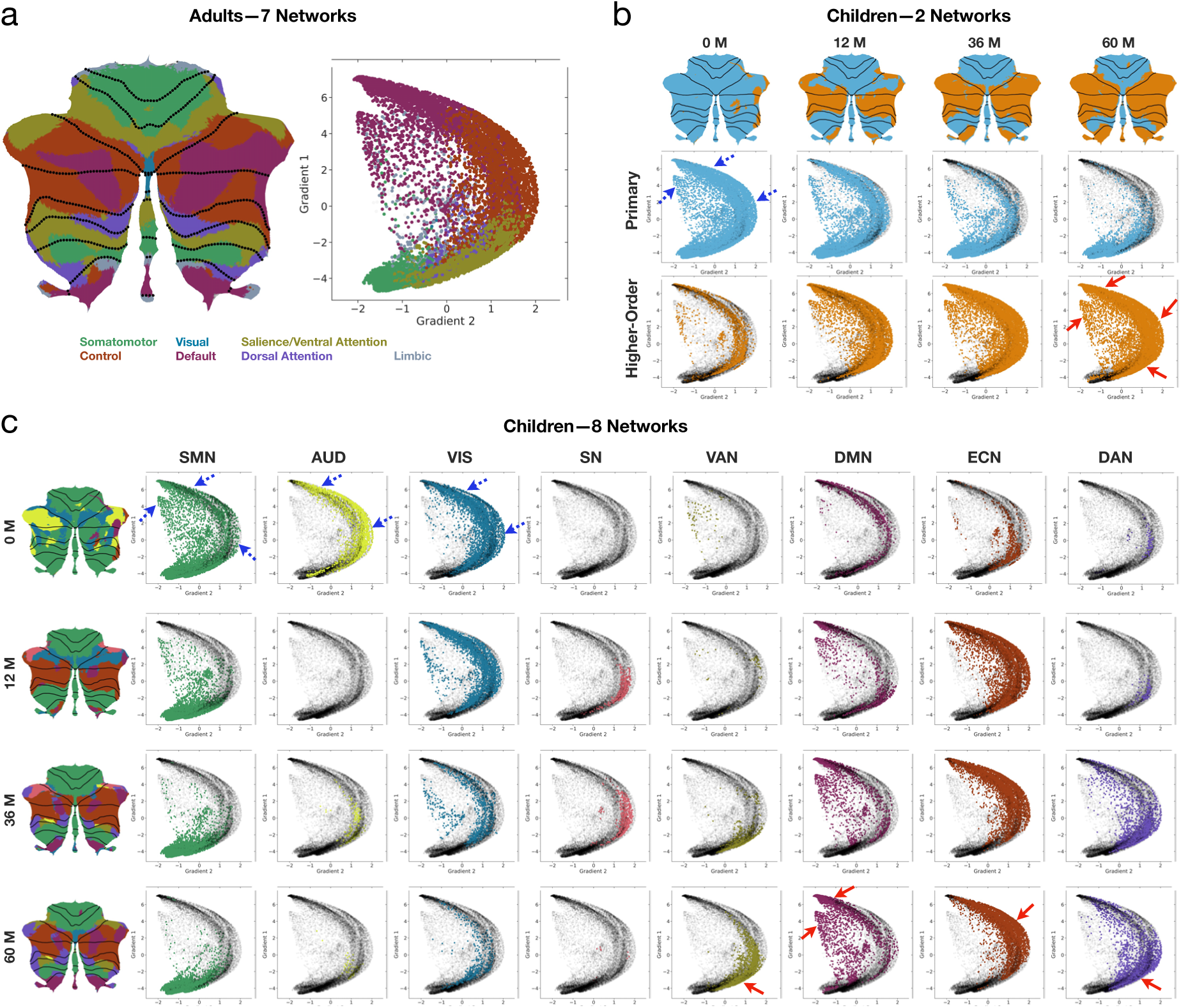
Functional gradients. **a**, Cerebellar functional gradients in adults^40, 42^. **b**, Functional gradients of primary and higher-order networks at birth, 12 months, 36 months, and 60 months. **c**, Network-specific functional gradients at birth, 12 months, 36 months, and 60 months. Gradient maps were generated using LittleBrain^42^, with each point representing a cerebellar voxel in the MNI space. Regions marked by blue dashed arrows on the gradient maps indicate areas where network points decrease or disappear with age, while regions marked by red solid arrows indicate areas where network points increase with age. The cerebellar gradient maps of large-scale networks spanning the entire early childhood period are shown in Figure S9.

We investigated how functional topography evolves over time by mapping our cerebellar parcellation maps onto the gradient map. At coarse granularity (Figure 5b), primary networks in early childhood initially cover nearly the entire map but gradually concentrate in regions with low Gradient 1 and low Gradient 2 as age increases. In contrast, higher-order networks, initially confined to a small region in middle Gradient 2, progressively expand into areas with high Gradient 1 and high Gradient 2 over time.

At medium granularity (Figure 5c and Figure S9), the gradient maps of different large-scale networks evolve differently with age:

- The SMN primarily varies along Gradient 1. With age, the presence of SMN in the high Gradient 1 diminishes. By 60 months, the SMN is predominantly concentrated in low Gradient 1 and low Gradient 2, in alignment with the SMN gradient pattern observed in adults.
- Although AUD initially occupies a large portion of high Gradient 1 and high Gradient 2 at birth, it declines thereafter. After six months, AUD appears only sporadically in maps and covers a limited area. Moreover, the number of points associated with the AUD and the VIS on the gradient maps reduces with age. Given the absence of AUD and the near-absence of VIS in the cerebellar gradient maps of adults, we speculate that the decline of AUD and VIS in children’s cerebellar gradient maps reflects a developmental reorganization of cerebellar connectivity, with decreasing direct involvement in auditory and visual processing and increasing engagement in higher-order cognitive functions. This shift may indicate a transformation in how the cerebellum integrates sensory information, potentially supporting more complex cognitive processes during early childhood.
- The DMN and ECN are present on the gradient map from birth and vary along Gradients 1 and 2, respectively. As age progresses, the distribution of the DMN becomes concentrated in high Gradient 1, while the ECN distribution becomes concentrated in high Gradient 2, resembling the adult-like pattern.
- Both the DAN and VAN occupy only small areas on the gradient map at birth. However, beginning at 24 and 36 months, respectively, the DAN and VAN gradually shift to the low Gradient 1 and slightly higher Gradient 2 regions, resembling their adult locations, and progressively expand over time.
- Since its emergence on the gradient map around 9 months, the SN is primarily concentrated in middle Gradient 1 and high Gradient 2, although its extent varies over time. Interestingly, this area intersects with the DMN, VAN, DAN, and ECN, underscoring the SN’s pivotal role in network switching^50, 51^.

Together, these gradient maps illustrate that the functional gradients of various large-scale networks evolve dynamically during early childhood. With development, most networks transition from a diffuse to a more focal distribution on the gradient maps, ultimately resembling adult-like patterns. Despite network-specific variations in cerebellar gradient maps during early childhood, peak concentration locations tend to stabilize around 36 months and align more closely with adult patterns. This stability highlights consistent functional organization across development, with core areas of network function and connectivity retaining fundamental patterns even as overall activity distributions become more refined.

### Temporal Trends

We charted temporal changes in cerebellar functional topography based on the volume fraction of each network determined based on winner-take-all parcellation (Figure 6a). At coarse granularity, although the proportion of cerebellar regions connected to primary networks is substantially larger than that of regions connected to higher-order networks at birth, it gradually decreases throughout early childhood. In contrast, the proportion of cerebellar regions connected to higher-order networks increases over time, gradually surpassing that of primary networks from approximately 24 months onward (Figure 6a, left column).

**Fig. 6.**
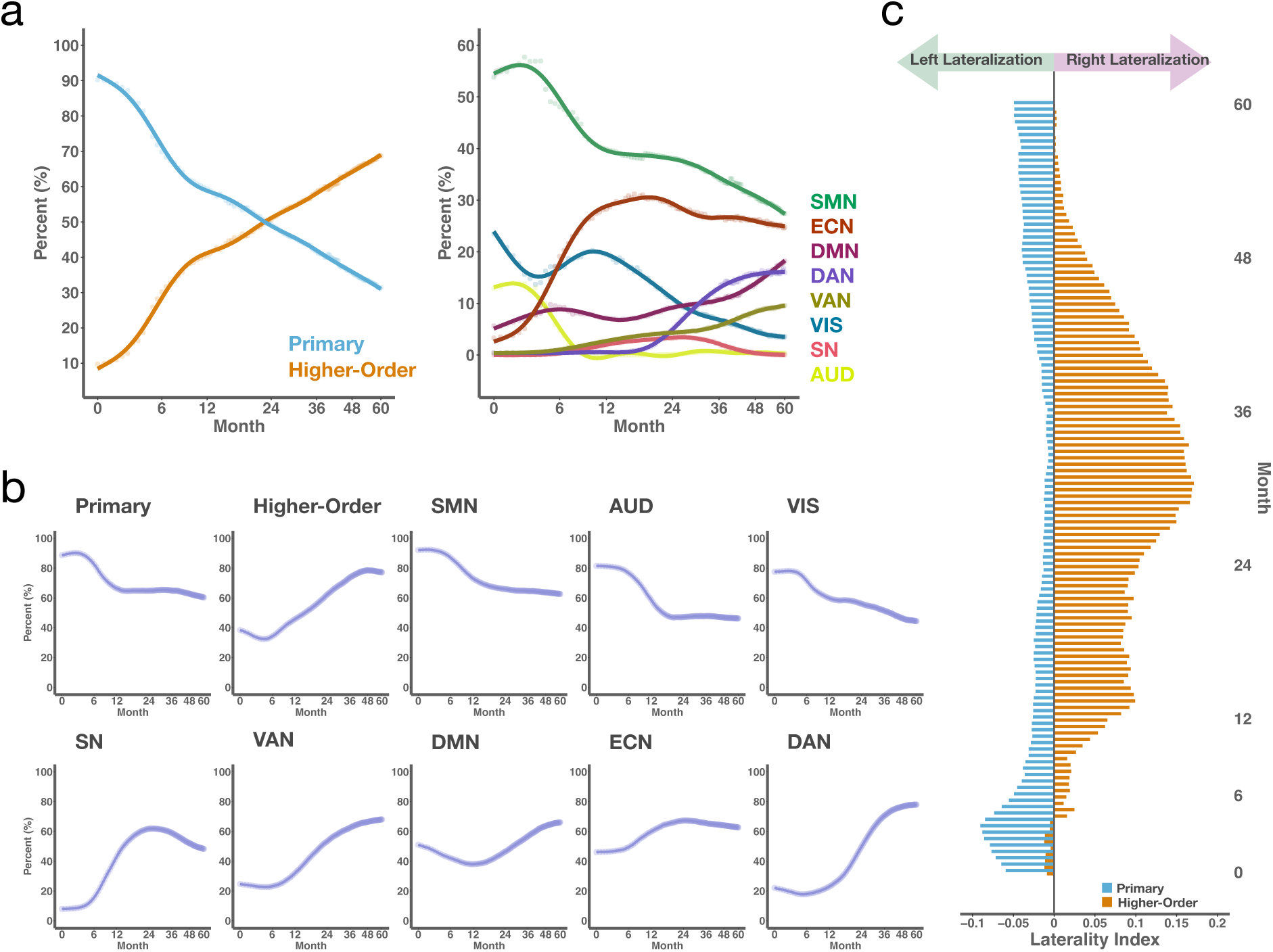
Developmental patterns of cerebellar functional topography. **a**, Trajectories of network-specific volume fractions at coarse granularity (left coloumn) and medium granularity (right coloumn).**b**, Trajectories of cerebellar volume fractions of voxels with positive connectivity to each cortical network during early childhood. **c**, Laterality of primary networks and higher-order networks from 0 to 60 months, with laterality index above 0 signifying right lateralization and below 0 signifying left lateralization.

At medium granularity, regions connected to the primary networks (SMN, AUD, and VIS) decrease, while those connected to higher-order networks (DMN, DAN, and VAN) generally increase throughout early childhood. An exception is the regions connected to the ECN, which expanded sharply until 12 months and then remained relatively stable, followed by a slight decline. However, cerebellar regions connected to the ECN have consistently been larger than those connected to other networks since 6 months, only smaller than those connected to the SMN. Notably, from birth to 6 months, most cerebellar regions are connected to the SMN, AUD, and VIS; from 6 to 24 months, the majority are connected to the SMN, ECN, and VIS; and from 24 to 60 months, the majority are connected to the SMN, ECN, and either the DMN or DAN (Figure 6a, right column). These developmental patterns suggest a dynamic shift in the cerebellum’s role from primarily sensory-motor processing to a gradual and increasing integration with higher-order networks, such as the ECN, DMN, and DAN, indicating a diversification of cerebellar functions during early childhood.

Given that the winner-take-all approach assigns each voxel exclusively to a single network, it may fail to capture overlapping connectivity patterns that reflect functionally relevant interactions. To overcome this limitation, we also calculated the volume fraction of cerebellar voxels that show positive connectivity with each network (Figure 6b). At coarse granularity, the volume fraction of primary networks generally declined from 0 to 12 months and remained relatively stable thereafter. In contrast, the volume fraction of higher-order networks showed a slight decline from 0 to 6 months, followed by a sustained increase. At medium granularity, the volume fractions of the SMN and AUD exhibited an overall decline from 0 to 18 months before stabilizing, whereas that of the VIS continuously decreased throughout early childhood. The volume fractions of the SN and ECN increased during the first 24 months but declined thereafter. The DMN’s volume fraction decreased during the first 12 months and then increased from 12 months onward, whereas the VAN and DAN exhibited a continuous increase throughout early childhood. Notably, during the first six months of life, changes in volume fractions based on positive connectivity were relatively small compared to those observed with the winner-take-all approach. This difference occurs because the winner-take-all method prioritizes the strongest connectivity, making it more sensitive to rapid local changes in network affiliations. This effect is especially pronounced during the first six months, a critical period of rapid cerebellocortical reorganization. Therefore, these two methods offer complementary perspectives on the development of cerebellocortical connectivity.

These results reveal dynamic changes in cerebellocortical functional connectivity during early childhood, showing how interactions between cerebellar regions and cortical networks evolve over time. This highlights the cerebellum’s adaptive role in processing sensorimotor and cognitive information, underscoring its influence on developmental trajectories.

### Functional Lateralization

The functional lateralization of primary motor and higher cognitive functions in the cerebellum is well established, with cerebro-cerebellar circuits considered to play a crucial role^52^. However, longitudinal research on cerebellocortical connectivity functional lateralization in developing children is currently lacking. To address this gap, we investigated cerebellar laterality with respect to the primary and higher-order networks from 0 to 60 months. We calculated the laterality indices by determining the number of significant connectivity voxels in the left and right cerebellar hemispheres connected to higher-order and primary networks at each age. We observed consistent leftward lateralization of the cerebellum’s significant connectivity with primary networks, as well as rightward lateralization of its significant connectivity with higher-order networks, which emerged after 4 months and remained stable throughout early childhood (Figure 6c), highlighting the distinct lateralization patterns between the cerebellum and these networks during early development. Our findings provide evidence, from the perspective of cerebellocortical functional connectivity, for functional changes resulting from unilateral cerebellar abnormalities in early life.

### Sex Differences

Sex differences in cerebellar gray and white matter volumes has been well-established in children and adolescents^53–55^. However, research has yet to explore sex differences in cerebellocortical functional connectivity during early childhood. To fill this gap, we investigated cerebellocortical functional connectivity separately in female and male children, aiming to elucidate sex-specific influences on these connections from early development. By examining cerebellocortical functional connectivity, we found that cerebellar regions exhibiting strong connections with each RSN are generally consistent between female (Figure S4) and male (Figure S5) children. To better illustrate sex differences during early childhood, we generated spatial difference maps (Figure S6), highlighting sex-specific variations in cerebellar functional connectivity for each RSN, and examined the temporal trajectories of peak cerebellar connectivity (Figure S7) in both sexes.

We generated cerebellar functional parcellation maps for female and male children (Figure 7a) to assess whether cerebellar functional topography exhibits sex-specific parttens. It can be observed from the coarse-granularity parcellation maps that the hierarchical organization of primary and higher-order networks emerged early in female children than in male children. The medium-granularity parcellation maps reveal that the Crus I and Crus II regions in female children are already prominently connected to the ECN and DMN at birth. In contrast, Crus I and Crus II in male children gradually become predominantly connected to the ECN and DMN around 9 months of age. Additionally, we generated cerebellar functional gradient maps for eight large-scale networks during early childhood based on the parcellation maps, separately for female (Figure S10) and male (Figure S11) children. As expected, the cerebellar functional gradients of female and male children show overall consistency with age, but sex-related variations persists for certain networks.

We further delineated the temporal trajectories of network volume fractions at both coarse and medium granularity levels for male and female children(Figure 7b). At coarse granularity, both sexes exhibit a higher proportion of cerebellar regions connected to primary networks than to higher-order networks at birth, with this difference being more pronounced in male neonates, but over time, the proportion of regions connected to higher-order networks increases while that of regions connected to primary networks decreases, so that by around 24 months, the proportion of cerebellar regions connected to higher-order networks surpasses that of primary networks. At medium granularity, the cerebellar regions connected to the SMN consistently occupy the largest proportion, regardless of sex. In female children, the proportion connected to the ECN consistently ranks second, except during the first month after birth. In male children, however, the proportion connected to the ECN does not reach second place until after 12 months; before then, the second-largest proportion is typically associated with the AUD or VIS. Our findings highlight both shared and divergent patterns of cerebellocortical connectivity in male and female children during early childhood and may contribute to research on sex-specific neurodevelopmental profiles.

**Fig. 7.**
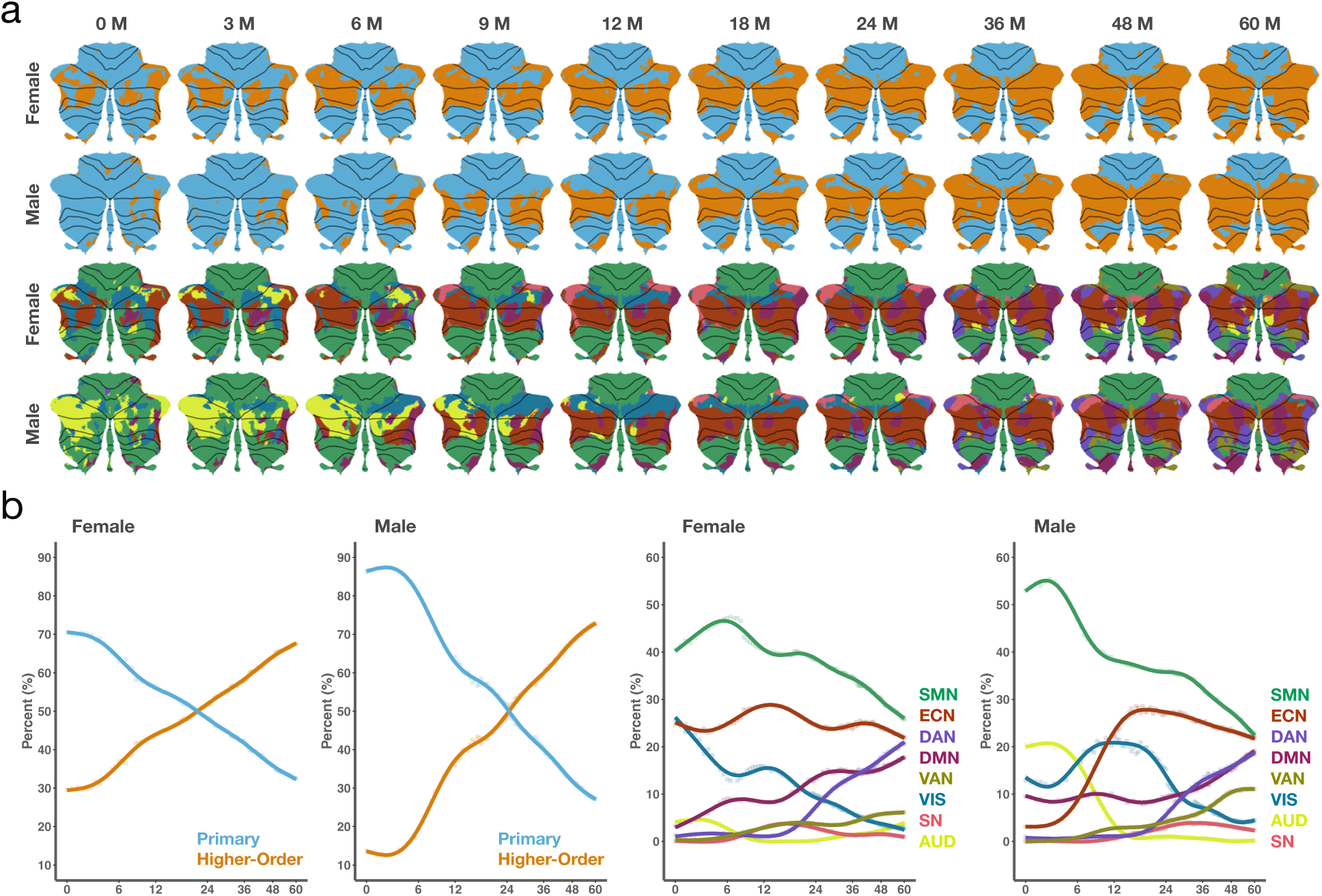
Spatiotemporal patterns of functional topography for female and male children. **a**, Parcellation maps for female and male children at coarse granularity (first two rows) and medium granularity (last two rows). **b**, Trajectories of network-specific volume fractions for female and male children at coarse granularity (left two columns) and medium granularity (right two columns).

## DISCUSSION

While extensive research has clarified the motor and cognitive functions of the cerebellum in adults^23, 40, 56, 57^, these roles remain poorly understood in early childhood—a critical period of rapid neural development. This stage involves ongoing synaptic pruning^18, 58^, which refines neural connections in response to development and stimuli, presenting a crucial window for neurodevelopmental intervention. Our study examined cerebellar connectivity with cortical networks and mapping its functional topography in children aged 0 to 60 months. Our findings show that cerebellar connectivity to higher-order networks is present at birth and generally strengthens throughout early childhood. We explored the hierarchical organization of cerebellocortical connectivity in early childhood and found that by around 36 months, cerebellar topography begins to resemble the adult pattern, becoming increasingly similar over time. We observed that the dispersed gradients of large-scale networks within the cerebellum during early childhood dynamically shift toward a more concentrated pattern with age, while generally maintaining stable anchor positions, similar to those observed in adults. We observed lateralized patterns of cerebellar connectivity, with higher-order networks being right-lateralized and primary networks being left-lateralized, further highlighting the cerebellum’s functional asymmetry. We identified male and female patterns in both the organization and development of cerebellocortical functional connectivity. These findings emphasize the early development of cerebellocortical functional connectivity, its role in neurodevelopment, and its potential influence on cognitive and motor outcomes, offering guidance for future research and clinical intervention in cerebellar dysfunction.

The cerebellum’s role in motor control and sensory modulation is well-established in humans^59^ and other species^60^, coordinating complex activities like skilled finger movements, eye positioning, and trunk and limb control. It is also crucial for motor learning^61^, essential for acquiring and refining motor skills. Damage to the cerebellum’s auditory and visual regions reduces accuracy in auditory- and visual-evoked orienting behaviors^60^, highlighting the cerebellum’s role in sensory processing. Cerebellar injury can impair visual target tracking^62^ and motion detection^63^, underscoring the cerebellum’s importance in sensory-motor integration. Despite these findings, few studies ^27, 64^ have investigated cerebellum-motor cortex functional connectivity in young children, and neuroimaging evidence of cerebellar connections with the AUD and VIS remains limited. Our study confirms that the developing cerebellum functionally connects with both the SMN and sensory regions such as the AUD and VIS. However, due to the lack of ground truth validation from optogenetics, histology, or tractography, these findings in early childhood remain to be fully confirmed. Notably, the proximity of the auditory cortex to the sensorimotor cortex has often led to its distinct cerebellar connections being overlooked. Recent research^65^ identified unique cerebellar functional connectivity with the auditory cortex, separate from its connectivity with the SMN. Our study adds to this understanding by showing that cerebellar functional connectivity with the AUD differs from the SMN not only in strength but also in developmental patterns, underscoring the cerebellum’s distinct role in auditory processing during early development. Moreover, we found strong cerebellar functional connectivity with the VIS in infants, in sharp contrast to the much weaker connectivity seen in adults, which is largely confined to the cerebellar vermis^26, 39^. This difference may reflect the early dominance of visual, auditory, and sensorimotor cortical hubs in childhood^66^.

While there is consensus on the existence of functional connectivity between the cerebellum and primary networks (at least motor networks), uncertainty remains regarding its functional connectivity with higher-order networks during early childhood (particularly in infancy). Research in this area is limited. While one study using diffusion tensor tractography suggests that structural connectivity between the cerebellum and higher-order cortical regions is established during infancy^67^, another functional MRI study found no evidence of cerebellar functional connectivity with these networks during this period^27^. This discrepancy highlights the complexity of cerebellocortical interactions in early childhood and underscores the necessity for further investigation. A recent study^68^ provides critical insight into these interactions, revealing that neonatal cerebellar injury can disrupt the development of higher-order cortical regions. Beyond motor impairments, neonates with isolated cerebellar injuries may exhibit cognitive, language, and social behavioral abnormalities during development^69, 70^, suggesting crucial functional connectivity between the cerebellum and higher cognitive networks during early development.

Infant learning was perceived as a passive accumulation of sensory experiences, predominantly associated with the sensory and motor cortex. However, emerging research ^71^ suggests that infants are intrinsically motivated for active learning, with the prefrontal cortex playing a key role in this process. Our study corroborates this view by revealing widespread cerebellar connectivity with higher-order networks during early childhood. Connectivity with higher-order networks is weaker than with primary networks but generally strengthens with age. Notably, specific higher-order networks, especially the ECN located in the frontal regions, exhibit robust connectivity with the cerebellum from birth. This finding is consistent with our previous research^72^, which highlighted strong structural connections between the cerebellum and the superior frontal gyrus—one of the key regions of the ECN. Our findings confirm that the functional connectivity between the cerebellum and higher-order networks is established early in life and may play a crucial role in shaping cognitive and behavioral functions during childhood, potentially influencing the developmental trajectory of these networks into adulthood. This early interaction may shed light on the underlying mechanisms of cognitive disorders such as ASD and ADHD, which are linked to abnormal cerebellar connections with higher-order networks^29–34^, offering new insights into how cerebellar development affects later cognitive functions.

The contralateral organization of cerebellar functional topography with the motor and prefrontal cortices is well-established in adults^19, 20^. Our findings extend this organization to early childhood, reinforcing the existence of the cross-hemispheric cerebrocerebellar loop^73^. Identifying this loop in early childhood may provide insights into contralateral developmental abnormalities in the cerebellum following unilateral cerebral damage in infancy, possibly due to remote trans-synaptic effects^8^. Beyond contralaterality, another key feature we observed is the hierarchical organization of primary and higher-order networks within the cerebellum, closely aligned with the S-A axis^45^. Extending from primary sensorimotor and visual regions to higher-order association areas^74, 75^, the S-A axis reflects a fundamental principle of hierarchical cortical organization, with spatiotemporal pattern of cortical maturation proceeding hierarchically along this axis^74, 76^. While the hierarchical organization along the S-A axis is well established for the cortex, corresponding research on the cerebellum remains limited. Guell et al.^24^ identified functional gradients in the cerebellum that mirror the cortical hierarchy, revealing a dual motor and triple nonmotor organization. However, unlike the S-A axis of cerebellocortical functional connectivity in adults^24, 77^, where gradient boundaries are near the superior posterior, prepyramidal, and secondary fissures, the first gradient boundary initially appears at the horizontal fissure after birth. This boundary shifts gradually, reaching the superior posterior fissure by around 24 months, while the second and third gradient boundaries resembling those in adults at the prepyramidal and secondary fissures. These findings indicate that the cerebellum’s hierarchical organization of primary and higher-order functions emerges early in infancy and progressively aligns with adult functional organization along the S-A axis. This alignment likely reflects the cerebellum’s early integration into broader cortical systems, offering critical insights into the cerebelloccortical functional connectivity patterns that support cognitive and motor development in early childhood.

In the past decade, research on the functional connectivity gradients of large-scale networks in the cerebral cortex has gained considerable attention^45, 78, 79^. A recent study^80^ mapped changes in cortical functional gradients across the entire human lifespan from cradle to grave. However, investigations on cerebellar functional gradients have predominantly focused on adults, represented primarily by the seminal work of Guell and colleagues^24, 25, 42^. The study of cerebellar gradients of large-scale networks in children, in particular, remain largely unexplored. For the first time, we mapped the functional gradients of large-scale networks in the cerebellum throughout early childhood, revealing how these gradients evolve during this critical period of brain development. In adults, distinct patterns of cerebellar gradients have been observed^24, 25, 42^: the motor network, ECN, and DMN occupy low, middle, and high positions on Gradient 1, respectively. The VAN and DAN are positioned along Gradient 1 away from the DMN, with the ECN situated between the VAN/DAN and DMN, acting as a mediator. Gradient 2 is thought to relate to task load, with low Gradient 2 corresponding to low task load networks such as the motor network and DMN, and high Gradient 2 to task-positive networks, including the VAN, DAN, and ECN. Not surprisingly, the cerebellar functional gradients at birth do not fully replicate the patterns observed in adulthood. Instead, these networks exhibit more dispersed gradients that gradually become more concentrated, converging towards an adult-like pattern as children mature. Since similar functions occupy adjacent gradient positions^78^, the progressive focalization of these gradients reflects functional specialization. Notably, despite dynamic changes in cerebellar gradients during early childhood, the anchor positions generally remain stable, mirroring those in adults. This suggests the foundational framework is likely established at birth and refined with age, potentially through synaptic pruning or activity-dependent synapse refinement, supporting the idea that early cerebellar organization shapes its mature functional architecture.

Studies on cerebellar functional asymmetry suggest that cognitive functions typically exhibit right laterality, while motor functions tend to show left laterality^52^. Consistent with this, our findings indicate that the cerebellum demonstrates rightward lateralization in connection with higher-order networks and leftward lateralization with primary networks throughout early childhood. Clinical studies^81, 82^ further support this pattern, revealing that damage to the right cerebellar hemisphere can lead to cognitive impairments, including deficits in language and literacy, whereas left cerebellar damage is frequently associated with specific visuospatial impairments and deficiencies in spatial skills. These findings highlight the role of lateralization in early neurodevelopment, showing that cerebellocortical functional connectivity supports both hemispheres, promoting balanced cognitive and motor development essential for effective interaction with the environment.

Although there are currently no reports on sex differences in cerebellocortical functional connectivity during development, it has been confirmed through cerebellar volumetric studies of infants, children, adolescents, and adults^53–55^. Our findings indicate that the cerebellum’s hierarchical organization develops later in male children than in females. Before the age of two, the proportion of cerebellar voxels predominantly connected to higher-order networks is greater in female than male children. This may contribute to a deeper understanding of the increased susceptibility of males to early-onset neuropsychiatric disorders^83^, such as ASD^31^ and ADHD^84^, which have been linked to reduced cerebellar connectivity with higher-order networks. In addition, our study indicates that female children exhibit noticeably stronger cerebellar connectivity with the SM-Tongue compared to male children during early childhood. This finding may provide neuroimaging evidence supporting earlier speech production^85^ and better verbal performance^86^ in female children. Our study highlights the importance of investigating sex differences in early cerebellocortical functional connectivity, as these differences may influence cognitive and behavioral outcomes. Additionally, it is crucial to recognize that factors such as genetics, nutrition, and socioeconomic conditions may also play a role in shaping functional connectivity^87^. Future research should account for these influences to better understand their contributions to cerebellocortical development and how they may interact with sex differences.

In summary, we investigated cerebellocortical functional connectivity during early childhood, spanning from birth to 60 months. Our findings demonstrate the presence of cerebellar connectivity not only with primary networks but also with higher-order networks, even at birth. Our parcellations with fine temporal resolution captured functional topography at different developmental stages, revealed the hierarchical organization of primary and higher-order networks, and suggested an S-A axis of early cerebellar functional development. Furthermore, our study shed light on the lateralization and sex-specific patterns of cerebellar functions. These findings offer insights into early neurodevelopment and may contribute to the diagnosis and monitoring of early-onset cerebellum-related disorders.

## METHODS

### Participants

The data utilized in this study were collected as part of the UNC/UMN Baby Connectome Project (BCP)^35^. After data preprocessing and quality control, the final dataset included 1,017 scans from 275 healthy participants (130 males and 145 females) ranging from birth to 60 months, with up to six longitudinal scans per participant. Parents were provided with full information about the study’s objectives before giving their written consent. Ethical approval for all study procedures was granted by the institutional review boards of the University of North Carolina at Chapel Hill (UNC) and the University of Minnesota (UMN).

### Data Acquisition

Children under 3 years old (0–35 months) were scanned while naturally asleep, without the use of sedatives. Prior to imaging, all babies were fed, swaddled, and fitted with ear protection. Children older than 3 years (36–60 months) were scanned while either asleep or watching a passive movie^35, 88^. All images were acquired using 3T Siemens Prisma MRI scanners (Siemens Healthineers, Erlangen, Gernamy) with 32-channel coils at the Biomedical Research Imaging Center (BRIC) at UNC and the Center for Magnetic Resonance Research (CMRR) at UMN^35^. MRI acquisition parameters are summarized as follows:

- T1 weighted MR images were obtained with a 3D magnetization prepared rapid gradient echo (MPRAGE) sequence: isotropic resolution = 0.8 mm, field of view (FOV) = 256 mm × 256 mm, matrix = 320 × 320, echo time (TE) = 2.24 ms, repetition time (TR)= 2400/1060 ms, flip angle = 8^◦^, and acquisition time = 6 min 38 s.
- T2 weighted MR images were obtained with a turbo spin echo (TSE) sequence: isotropic resolution = 0.8 mm, FOV = 256 mm × 256 mm, matrix = 320 × 320, TE = 564 ms, TR = 3200 ms, and acquisition time = 5 min 57 s.
- Resting-state functional MRI data were collected using a single-shot echo-planar imaging (EPI) sequence: isotropic resolution = 2 mm, FOV = 208 mm × 208 mm, matrix =104 × 104, TE = 37 ms, TR = 800 ms, flip angle = 52^◦^, and acquisition time = 5 min 47 s.

### Data Preprocessing

fMRI data processing is summarized in Figure S1. The fMRI blood oxygen level dependent (BOLD) data were first minimally preprocessed as follows: (i) Head motion correction using FSL `mcflirt` function^89^, rigidly registering each fMRI time frame to a single-band reference (SBref) image and generating the corresponding motion parameter files; (ii) EPI distortion correction using FSL `topup` function^90, 91^, generating a distortion correction deformation field using a pair of reversed phase-encoded FieldMaps; (iii) Rigid registration (degrees of freedom = 6) of SBref to FieldMaps; (iv) Rigid boundary-based registration (BBR)^92^ of distortion corrected SBref image to the corresponding T1-weighted (T1w) images, with prealignment using mutual information as the cost function; and (v) One step sampling using combined deformation fields and translation matrices, resulting in motion- and distortion-corrected fMRI data in the subject native space (T1w space)^93^.

### Data Denoising

We denoised the minimally preprocessed fMRI data before further analysis. We detrended the data to remove slow drift by applying a high pass filter with a cutoff frequency of 0.001 Hz. We then utilized Independent Component Analysis-based Automatic Removal Of Motion Artifacts (ICA-AROMA)^94, 95^ to mitigate any residual motion artifacts. This process involves performing a 150-component independent component analysis (ICA) on the fMRI data and classifying each component as either BOLD signal or artifact based on high-frequency contents, correlation with realignment parameters (i.e., motion parameters estimated in head motion correction), edges, and CSF fractions. Components identified as motion-related artifacts are non-aggressive removed through regression. We subsequently mapped the denoised fMRI data to the MNI space^96, 97^. We then applied Gaussian smoothing with a full-width at half-maximum (FWHM) of 4 mm separately to the cerebrum and cerebellum. The cerebral and cerebellar masks were obtained by segmenting the high-resolution MNI-ICBM 152 symmetrical template^96–98^ and downsampled to a 2 mm resolution. Finally, we intensity-normalized the data to a constant mean volume intensity of 10,000^99^, reducing scanner-related intensity variations and enabling reliable cross-subject comparisons for subsequent functional connectivity analyses.

### Quality Control

To ensure good data quality, we computed framewise displacement (FD) from motion parameters obtained through head motion correction. Samples with mean Power’s FD (absolute sum of motion parameters) exceeding 0.5 mm—a common threshold used in fMRI studies^100^—were excluded. Additionally, we set a limit for Jenkinson’s FD^89^ (Euclidean sum of motion parameters) at 0.2 mm to further minimize the impact of motion-affected samples. Out of 1,656 preprocessed fMRI samples, 1,493 met the Power’s FD criterion, and 1,278 further satisfied the Jenkinson’s FD criterion. We then visually evaluated the quality of skull stripping, tissue segmentation, and fMRI-to-T1w image registration, retaining 1,245 with satisfactory quality. The fMRI data in MNI space were categorized based on geometric distortions as “pass” (no or light distortion), “questionable” (light to medium distortion), and “fail” (medium to large distortion). After visual inspection, 1,063 samples were labeled as having “questionable” or better quality, out of which 703 were labeled as “pass”. Samples with “fail” quality were excluded from our analysis. We judiciously used the data according to these categories. We computed template spatial maps using only the “pass” labeled data to minimze artifacts in determining meaningful functional networks. We eliminated 46 from the 1,063 initially labeled as “pass” or “questionable” due to erroneous results in partial correlation analysis, resulting in a final total of 1,017 samples for subsequent analyses.

### Independent Component Analysis and Network Identification

We performed probabilistic group independent component analysis (GICA) exclusively on the “pass” fMRI data to ensure fewer noise-related components, ultimately yielding 40 RSNs, which served as templates for subsequent analyses. An anatomical examination of the template RSNs revealed that 30 were located in the cortex, while 10 were in the cerebellum or subcortical areas. We further categorized the 30 cortical RSNs into eight large-scale networks^20, 101^ (Table S1): SMN, AUD, VIS, SN, VAN, DMN, ECN, and DAN. RSNs within the SMN were named according to major motor functions, while those in other large-scale networks were named based on their anatomical locations. The terms “Left” and “Right” designate the hemisphere (left or right) in which each component is situated.

### Individual RSNs via Dual Regression

Using the template RSNs, we computed the RSNs for each individual based on their denoised fMRI data via dual regression^102^. We then applied Gaussian mixture modeling to each individual RSN to create activation probability maps. These maps were used to generate personalized cerebral masks for computing cerebellocortical functional connectivity in subsequent analyses.

### Cerebellocortical Functional Connectivity

Cerebellocortical functional connectivity (FC) was computed for each voxel in the cerebellum relative to a set of cortical functional networks. First, we applied a Butterworth bandpass filter (0.008–0.1 Hz) to the BOLD time series^103, 104^. To avoid cortical voxel influence from nearby cerebellar voxels, we expanded the cerebellar mask by 8 × 8 × 8 mm^3^ and excluded cortical voxels within it. For each of the 30 cortical RSNs, we extracted a representative time series by computing the major eigen time series of the denoised BOLD signals from the top 10% most activated voxels within the cerebral mask^19^. The eigen time series is the first eigenvector from singular value decomposition (SVD), with its sign adjusted^105^ to match the mean voxel time series. FC was quantified as partial correlation scores between the cortical eigen time series and the denoised BOLD time series of each cerebellar voxel within the non-dilated cerebellar mask. The correlation scores were Fisher’s r-to-z transformed, followed by standardization to unit variance.

### Spatiotemporal Maps

We used a generalized additive mixed model (GAMM) to analyze cerebellocortical FC at each voxel, implemented with the `gamm4` package in R^106^. The model included a smooth age term and random effects for subject ID and scan site: *Y* ∼ *s*(age, *k* = 10) + (1|SubjectID) + (1|SiteID), where *s*(·), where *s*(·) represents the smooth term and (1|·) models random effects. The number of basis functions, *k*, was empirically set to slightly exceed the degrees of freedom of the data. Three separate GAMMs were fitted for males, females, and both sexes, generating FC saptial maps from birth to 60 months at half-month intervals for a total of 121 time points. Gaussian mixture modeling in FSL MELODIC was applied to the predicted FC across time points to estimate the mean and variance of the null distribution, which was then normalized to have zero mean and unit variance. Peak FC was quantified as the 99th percentile of the FC spatial map. Spatial maps of sex differences were obtained by subtracting male FC maps from female FC maps and standardizing to unit null distribution variance. Medium- and coarse-granularity spatial maps (Table S1) were obtained by taking the signed absolute extrema of the fine-granularity maps.

### Parcellation

We generated parcellation maps from the 30 cerebellocortical FC spatial maps using a winner-take-all strategy. First, we applied 7 mm FWHM spatial smoothing to each FC map to reduce spatial noise. Next, we used probabilistic threshold-free cluster enhancement (pTFCE)^107^ to generate enhanced p-value maps, which were then upsampled to a resolution of 0.5 × 0.5 × 0.5 mm^3^. A winner-take-all approach assigned each cerebellar voxel to the FC component with the lowest enhanced p-value, creating cerebellar functional parcellations for each time point. Using this method, we generated fine-granularity parcellation maps, which were then grouped according to Table S1 to produce medium- and coarse-granularity maps. Volume fraction trajectories, derived from the parcellation, were modeled using thin-plate splines via the `gam` function in R: `volume_fraction` ∼ *s*(`age`), with 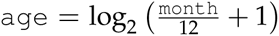. A logarithmic time scale was used to capture rapid changes during the first year of life.

### Functional Laterality

We quantified functional laterality based on the volume fractions of voxels in the right and left cerebellar hemispheres with connectivity greater than 3. The laterality index (LI) was calculated as 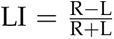, where *R* and *L* represent the volume fractions in the right and left cerebellar hemispheres, respectively. The LI ranges from −1 (total leftward lateralization) to 1 (total rightward lateralization), with 0 indicating complete symmetry.

### Effects of Wakefulness and Motion

We assessed the impact of wakefulness in children over 3 years old and motion artifacts on our results by testing whether unexplained variance in the GAMM residuals could be attributed to these factors. To test this, we used the following generalized additive model (GAM): `residuals` ∼ `wakefulness` + `motion`, where `residuals` are GAMM residuals after removing random effects, `wakefulness` is a binary variable indicating whether the subject is older than 3 years, and `motion` is a continuous variable representing the mean frame displacement of a fMRI scan ^100^. Table S2 reports the percentage of voxels with significant `wakefulness` and `motion` effects.

### Functional Correlation with Mullen Scores

We investigated the relationship between cerebellocortical FC and scores from the Mullen Scales of Early Learning (MSEL)^108^, using the following steps: (i) Normalize invididual cerebellocortical FC spatial maps to have a null distribution of zero mean and unit variance. (ii) Spatially smooth (7 mm FWHM) spatial maps to reduce noise. (iii) Consolidate medium- and coarse-granularity spatial maps using signed absolute extrema. (iv) Extract the peak FC of each spatial map. (v) Adjust peak FC and Mullen scores for age effects with a linear mixed model (LMM): *Y* ∼ age + (1|subject ID). (vi) Compute Pearson correlations between LMM residuals of peak FC and Mullen scores. The results are shown in Table S3.

### Visualization

Cortical functional networks were visualized using MRIcroGL (https://www.nitrc.org/projects/mricrogl/). Cerebellocortical FC flat maps were displayed using the Statistical Parametric Mapping software package (SPM12, https://www.fil.ion.ucl.ac.uk/spm/software/spm12/) and the SUIT^36^ software package (https://github.com/jdiedrichsen/suit). Functional gradient maps were generated with LittleBrain^42^. Trajectories of cerebellocortical FC across age were analyzed in R and visualized using `ggplot2`^109^.

## Supplementary Information

The manuscript contains supplementary material.

## Acknowledgements

The authors thank Jörn Diedrichsen, Caroline Nettekoven, and Aikaterina Maroli for their valuable discussions. This work was supported in part by the United States National Institutes of Health (NIH) under grants R01 MH125479, R01 EB008374, R01 EB035160, and R01 MH133836.

## Author Contributions

W.L.: methodology, investigation, visualization, data curation, writing – original draft, writing — review and editing. K.-H.T.: data curation, methodology, software, visualization, writing — review and editing. K.M.H.: resources, writing — review and editing. L.W.: resources. W.L.: resources. S.A.: resources, writing — review and editing. P.-T.Y.: conceptualization, supervision, funding acquisition, investigation, validation, writing — review and editing.

## Data Availability

The data used in this work is available via the National Institute of Mental Health data archive (NDA, https://nda.nih.gov).

## Code Availability

Software packages used in this work include FSL v.6.0.7.6, ICA-AROMA, ANTs, MATLAB (toolboxes include BrainNet Viewer, Network Community Toolbox, SPM12, SUIT, etc.), Python v.3.12, R v.3.8.0 (packages include gamm4, mgcv, bigmemory, ggplot2, etc.), and MRIcroGL.

---

**Fig. S1.**
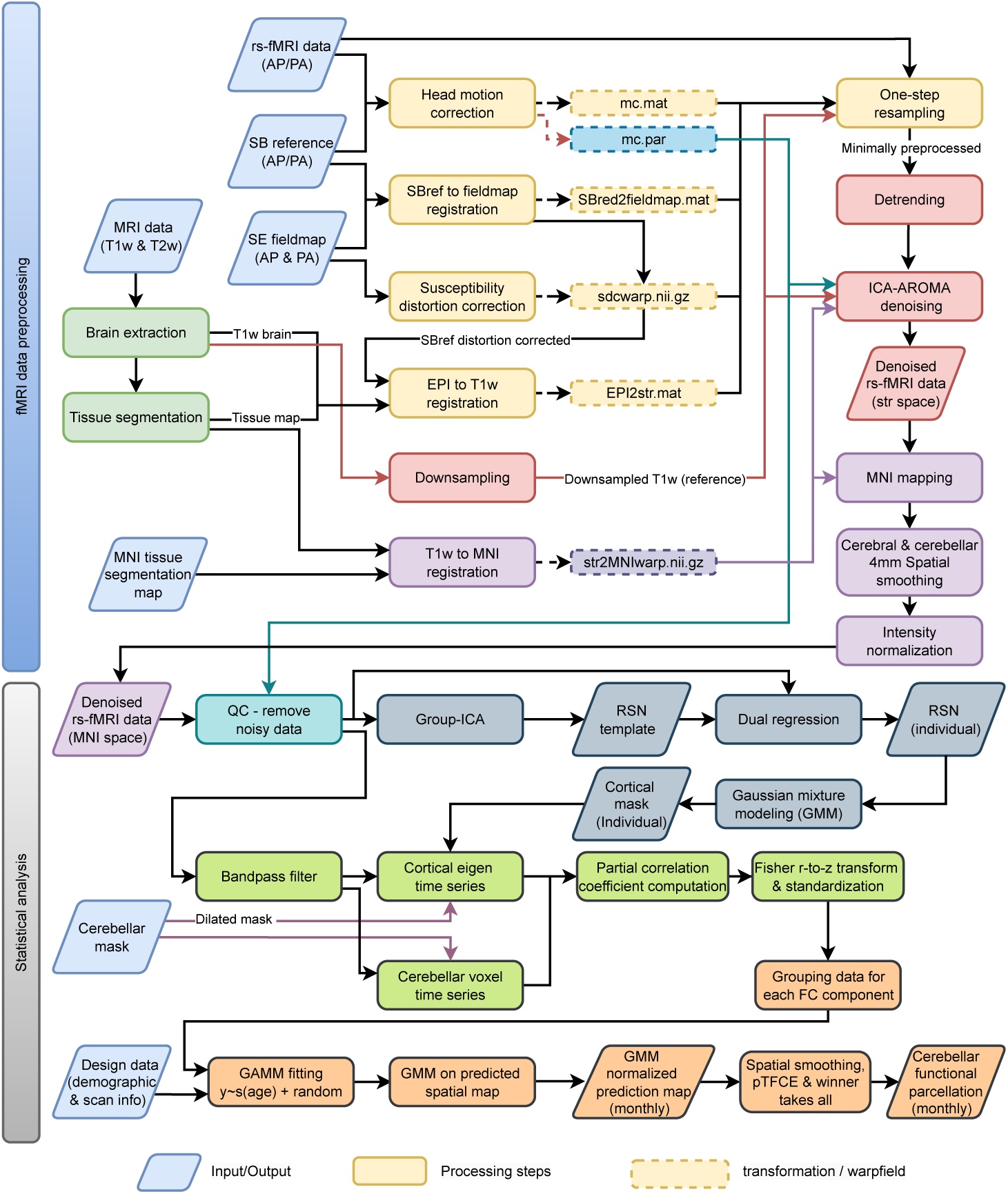
Processing and analysis pipelines. Steps involved in data processing and statistical analysis.

**Tab. S1.** Networks. Cortical networks and their corresponding primary regions.

| Network | Abbreviation | Primary Regions |
| --- | --- | --- |
| **Sensorimotor Network (SMN)** |  |  |
| Foot | SM-Foot | Bilateral precentral and postcentral gyri |
| Left hand | SM-Hand-L | Left precentral and postcentral gyri |
| Right hand | SM-Hand-R | Right precentral and postcentral gyri |
| Tongue | SM-Tongue | Bilateral precentral and postcentral gyri |
| **Auditory Network (AUD)** |  |  |
| Auditory | AUD | Bilateral superior temporal gyri |
| **Visual Network (VIS)** |  |  |
| Medial occipital | VIS-Occ-Med | Bilateral cuneus and lingual gyri |
| Lateral | VIS-Lat | Bilateral cuneus, lingual gyri, and fusiform gyri |
| Superior occipital | VIS-Occ-Sup | Bilateral cuneus |
| Inferior occipital | VIS-Occ-Inf | Bilateral cuneus and lingual gyri |
| Occipital pole | VIS-Occ-Pol | Bilateral occipital poles |
| **Salience Network (SN)** |  |  |
| Medial | SAL-Med | Anterior cingulate cortex and bilateral insulae |
| Lateral | SAL-Lat | Bilateral inferior frontal gyri and insulae |
| **Ventral Attention Network (VAN)** |  |  |
| Frontal | VA-Front | Bilateral superior, middle, and inferior frontal gyri |
| Parietal | VA-Par | Bilateral paracentral lobes, supramarginal gyri, and insulae |
| **Default Mode Network (DMN)** |  |  |
| Prefrontal | DM-Pref | Prefrontal cortex |
| Posterior cingulate | DM-Cing-Post | Posterior cingulate cortex and precuneus |
| Parietal angular | DM-Ang | Bilateral angular and middle temporal gyri |
| Left temporal | DM-Temp-L | Left middle temporal gyrus and temporal pole |
| Right temporal | DM-Temp-R | Right middle temporal gyrus and temporal pole |
| Parahippocampal | DM-PHippo | Bilateral parahippocampal, fusiform, and angular gyri |
| **Executive Control Network (ECN)** |  |  |
| Prefrontal | EC-Pref | Prefrontal cortex |
| Left frontal | EC-Front-L | Left superior and middle frontal gyri |
| Right frontal | EC-Front-R | Right superior and middle frontal gyri |
| Supramarginal | EC-SMarg | Bilateral supramarginal gyri |
| Temporal | EC-Temp | Bilateral fusiform and inferior temporal gyri |
| **Dorsal Attention Network (DAN)** |  |  |
| Medial parietal | DA-Par-Med | Bilateral precuneus and prefrontal cortices |
| Left parietal | DA-Par-L | Left superior parietal lobule |
| Right parietal | DA-Par-R | Right superior parietal lobule |
| Supramarginal | DA-SMarg | Bilateral supramarginal and inferior frontal gyri |
| Temporal | DA-Temp | Bilateral angular and inferior temporal gyri |

**Fig. S2.**
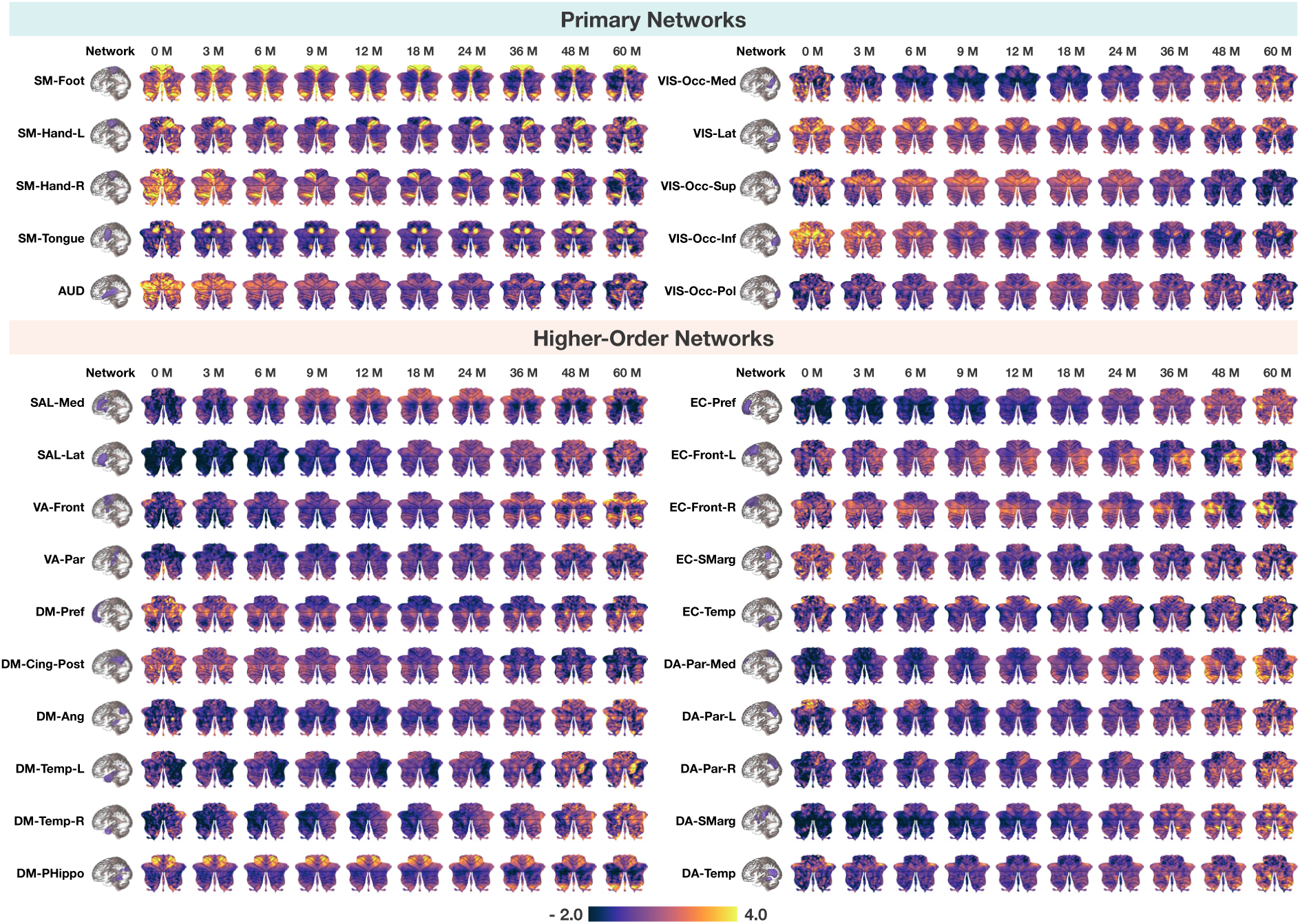
Cerebellar functional maps from birth to 60 months for children of both sexes. Spatiotemporal patterns of cerebellocortical functional connectivity (z-transformed) between the cerebellum and each RSN across early childhood. Values outside the range of −2.0 to 4.0 are capped for clarity.

**Fig. S3.**
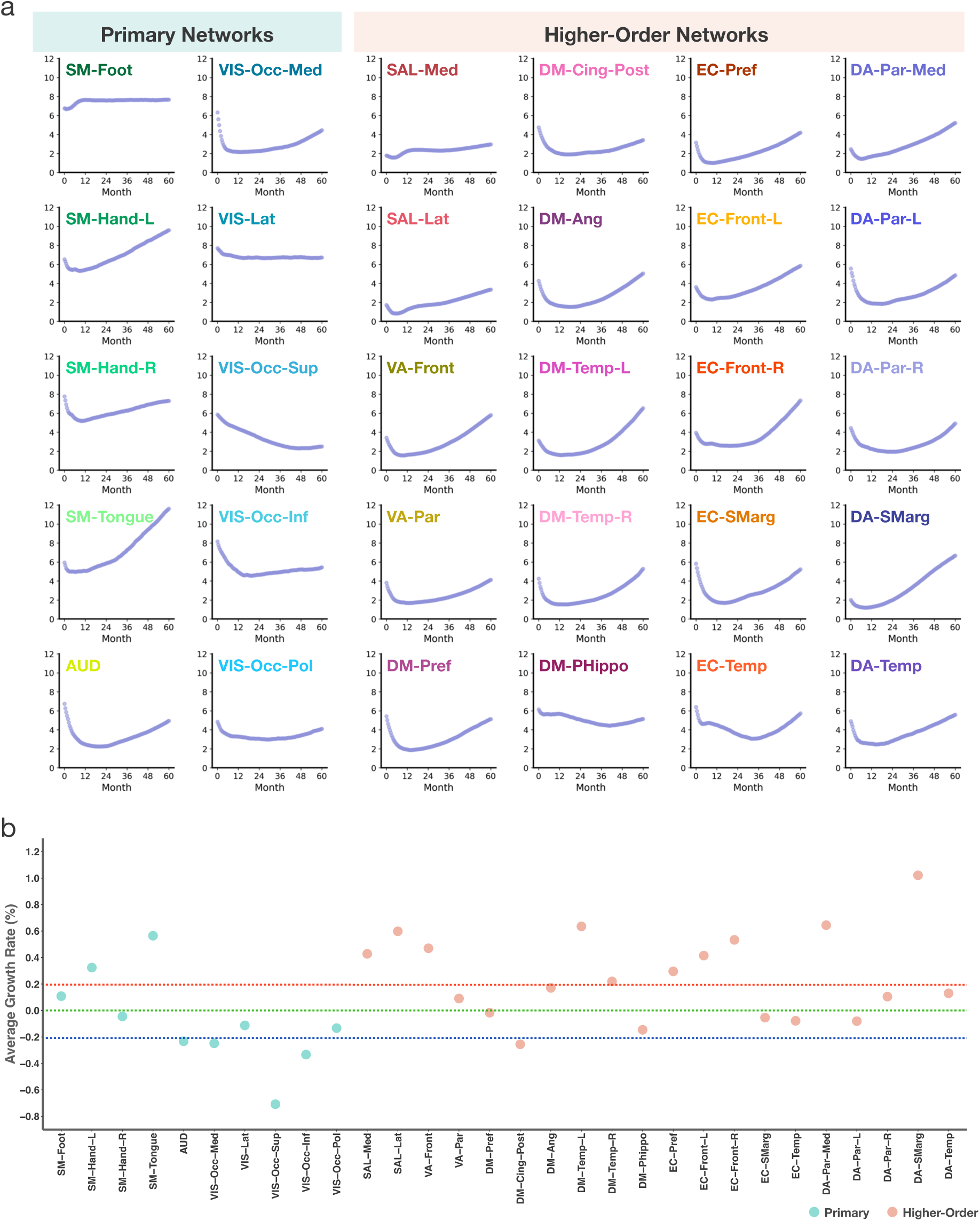
Developmental trends of cerebellocortical functional connectivity. **a**, Trajectories of the peak cerebellar connectivity (z-transformed) with each RSN over time. **b**, Average biweekly growth rate of cerebellar connectivity with each RSN. Values above the red dashed line denote substantial positive growth, values around the green dashed line denote negligible or minimal growth, and values below the blue dashed line denote substantial negative growth.

**Fig. S4.**
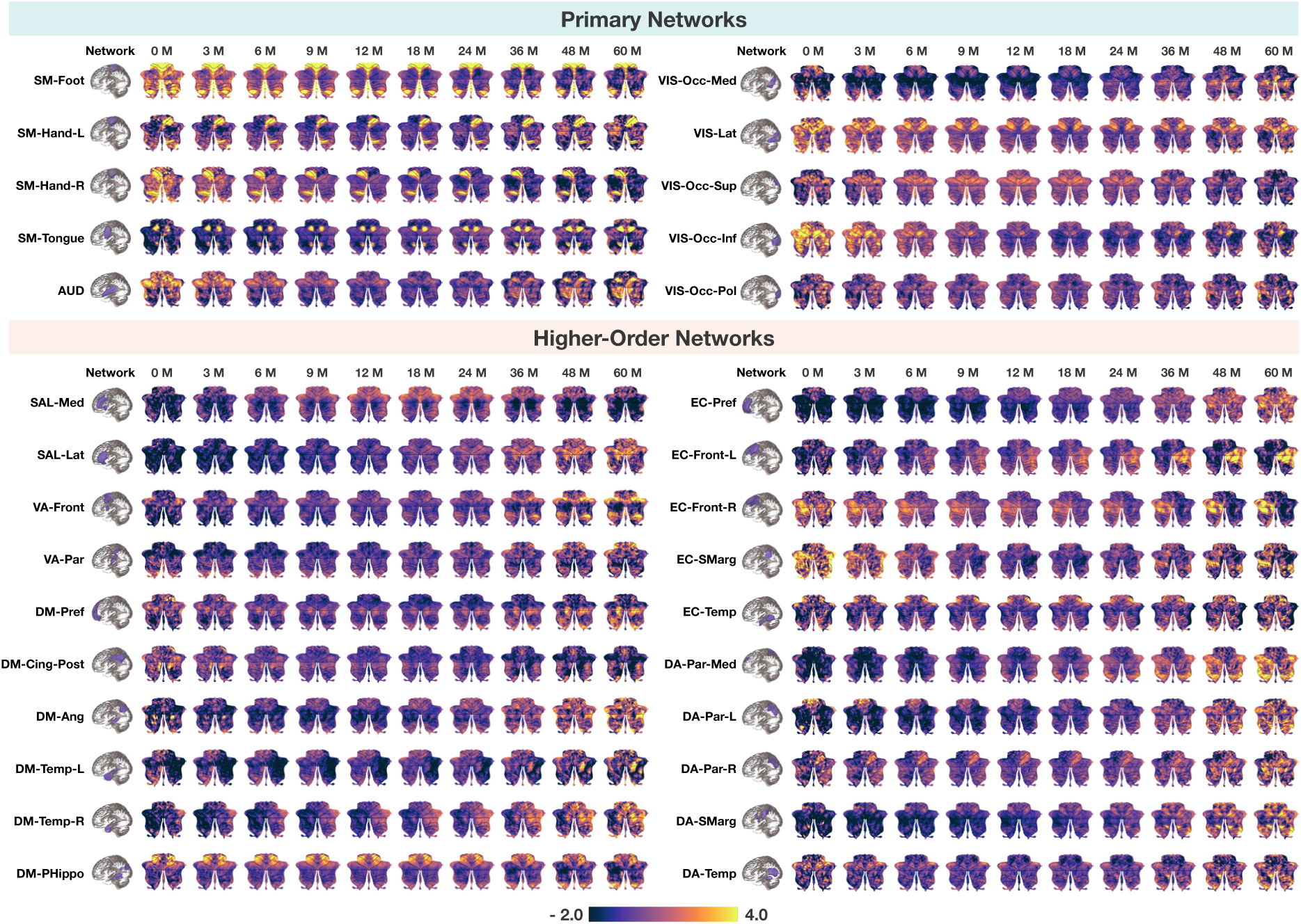
Cerebellar functional maps from birth to 60 months for female children. Spatiotemporal patterns of cerebellocortical functional connectivity (z-transformed) between the cerebellum and each RSN during early childhood in females. Values outside the range of −2.0 to 4.0 are capped for clarity.

**Fig. S5.**
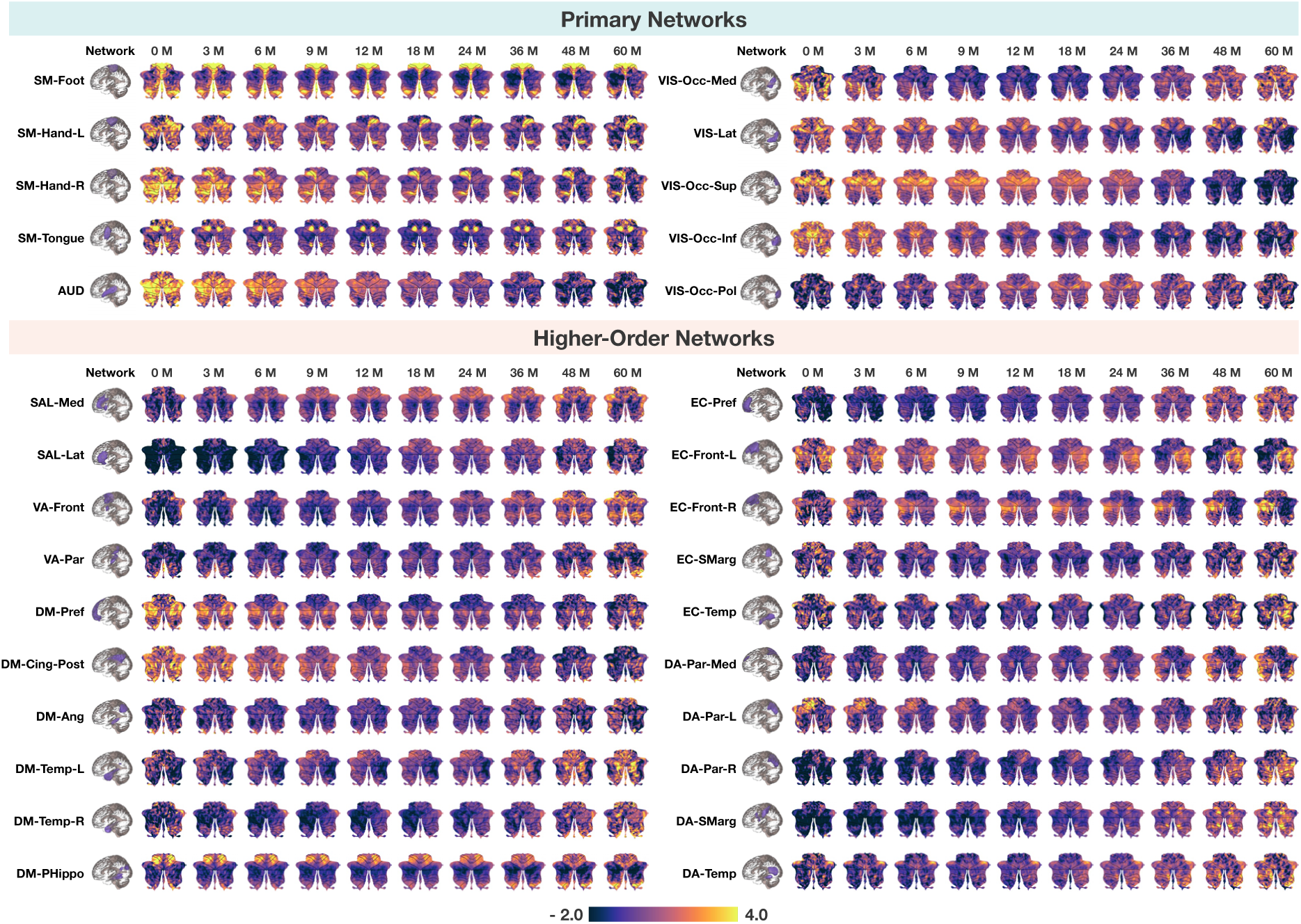
Cerebellar functional maps from birth to 60 months for male children. Spatiotemporal patterns of cerebellocortical functional connectivity (z-transformed) between the cerebellum and each RSN during early childhood in males. Values outside the range of −2.0 to 4.0 are capped for clarity.

**Fig. S6.**
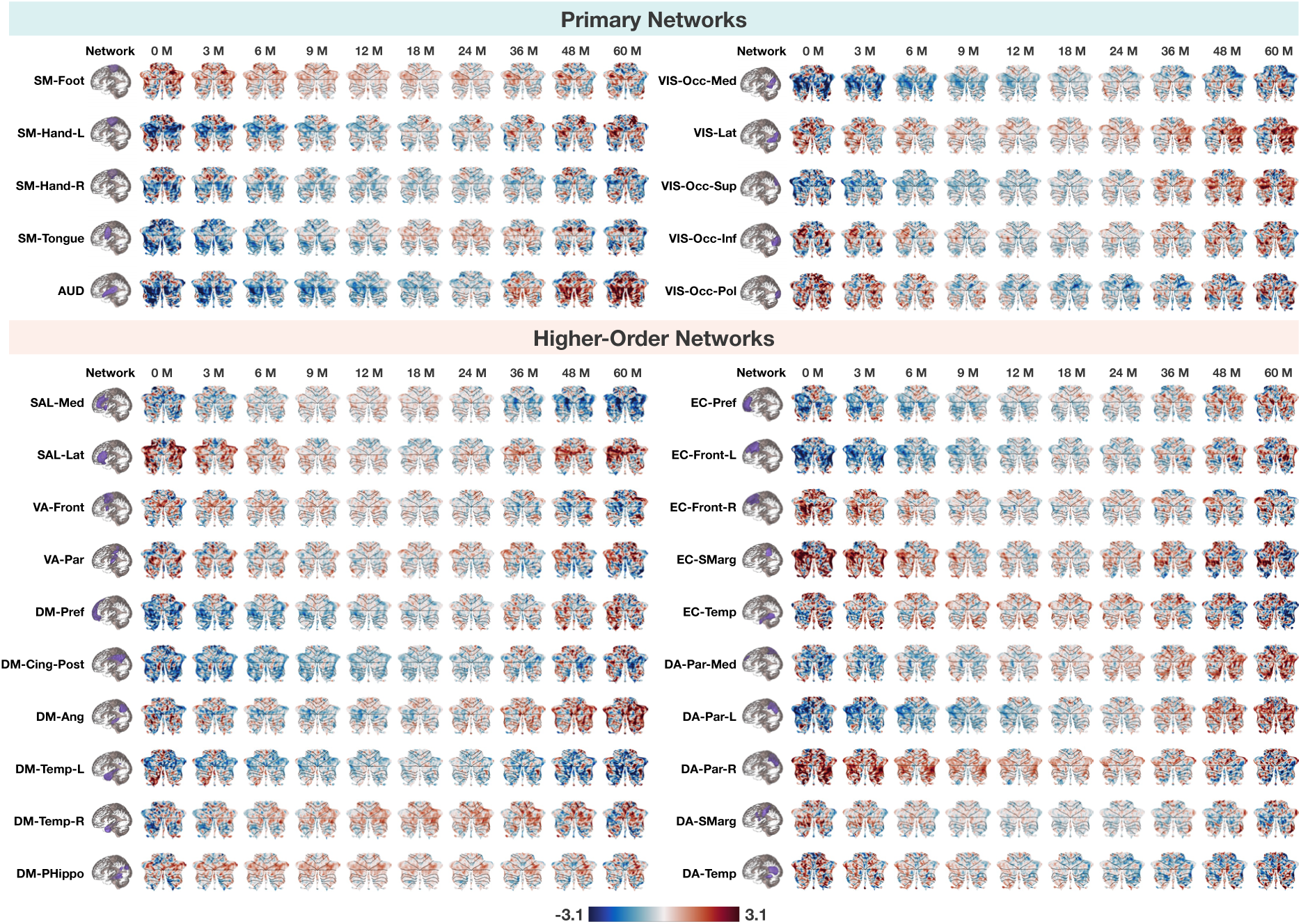
Sex differences spatial maps in cerebellocortical functional connectivity from birth to 60 months. Spatiotemporal patterns of sex differences in cerebellocortical functional connectivity with each RSN during early childhood are shown. For clarity, values exceeding ±3.1 (*p <* 0.001) are capped. The color scale represents the direction and magnitude of the differences, with red indicating stronger connectivity in females and blue indicating stronger connectivity in males.

**Fig. S7.**
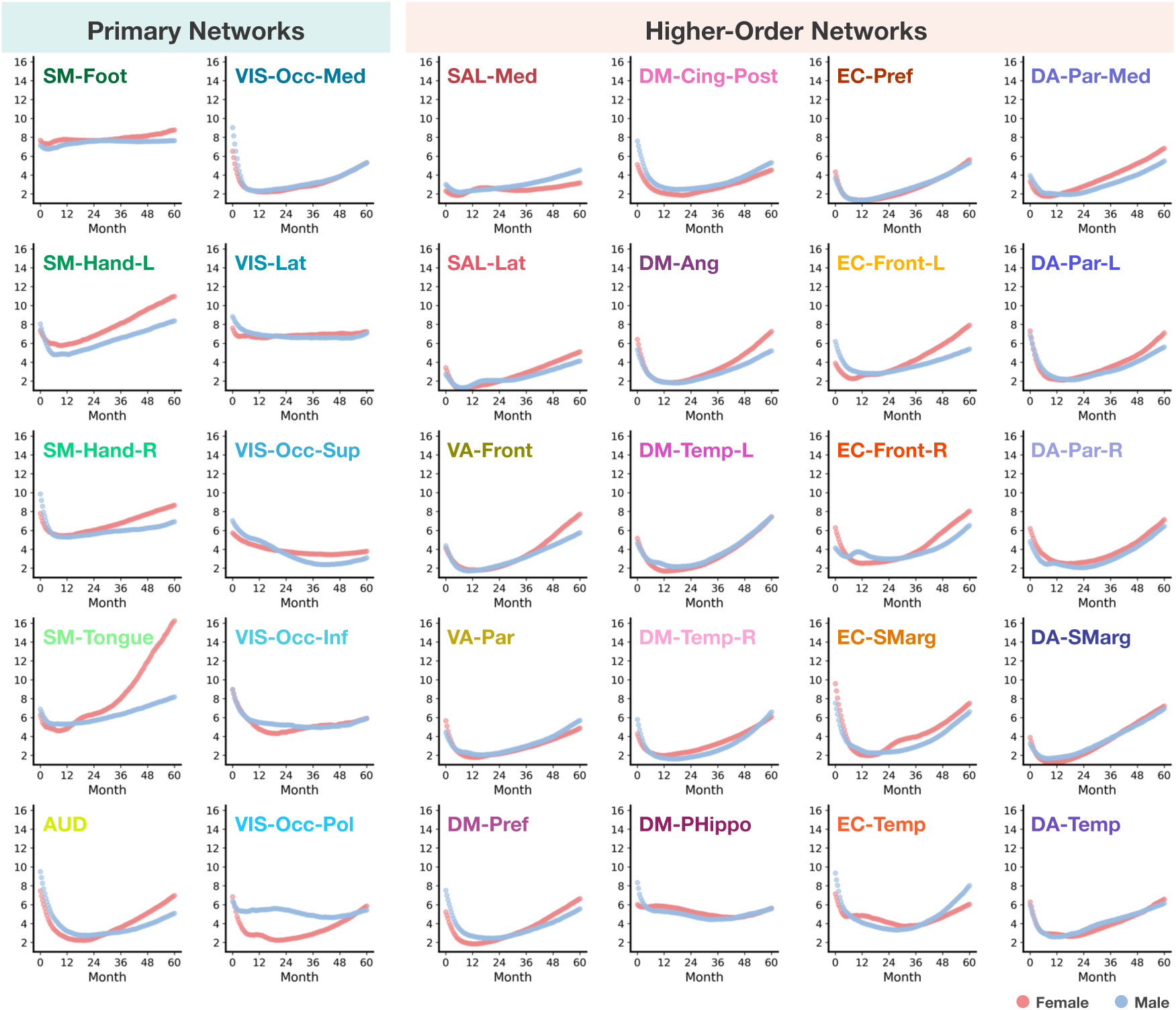
Developmental trends of cerebellocortical functional connectivity in female and male children. Trajectories of the peak cerebellar connectivity (z-transformed) with each RSN over time across female and male children.

**Fig. S8.**
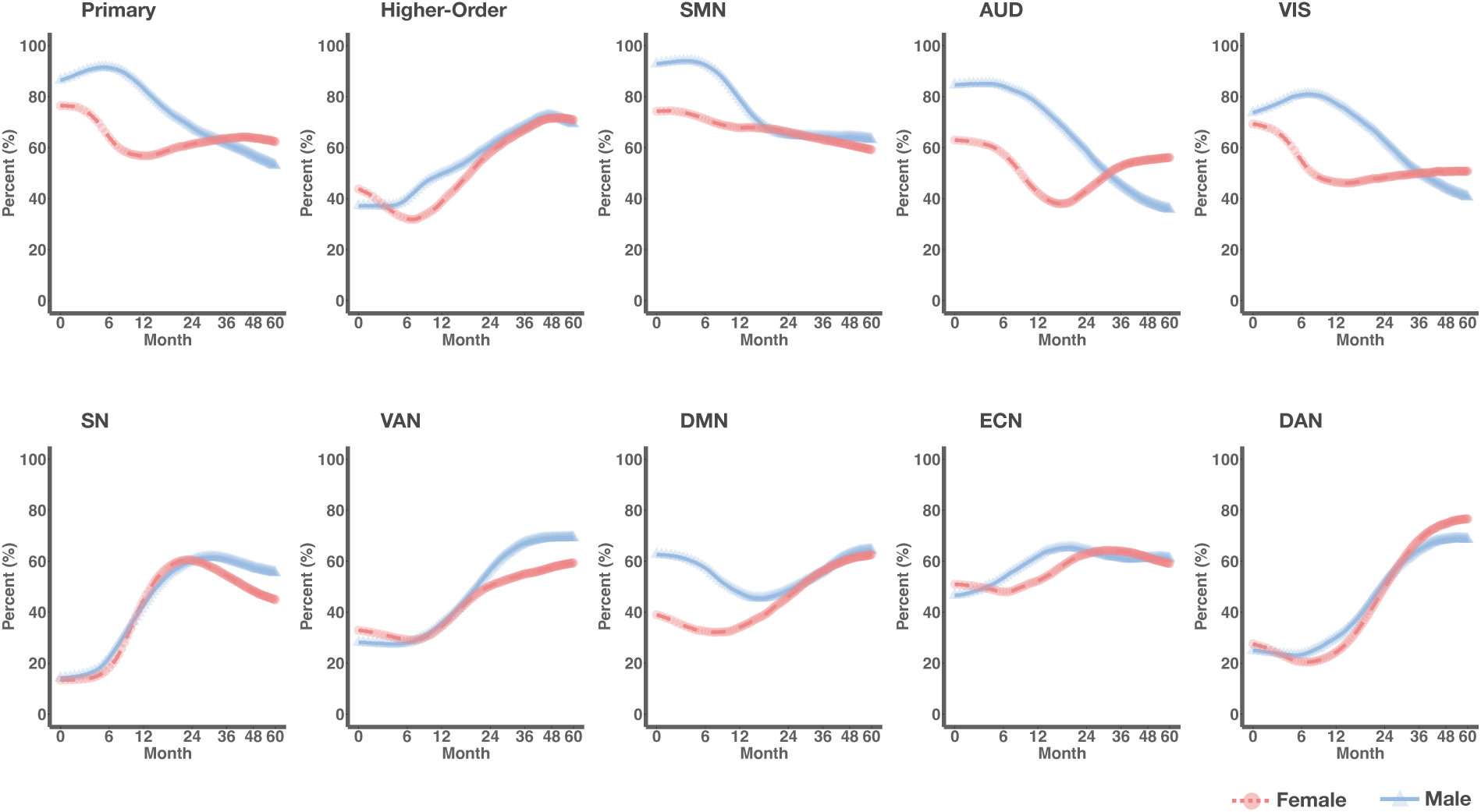
Sex-specific trajectories. Trajectories of cerebellar volume fractions of voxels with positive connectivity to cortical networks in female and male children.

**Fig. S9.**
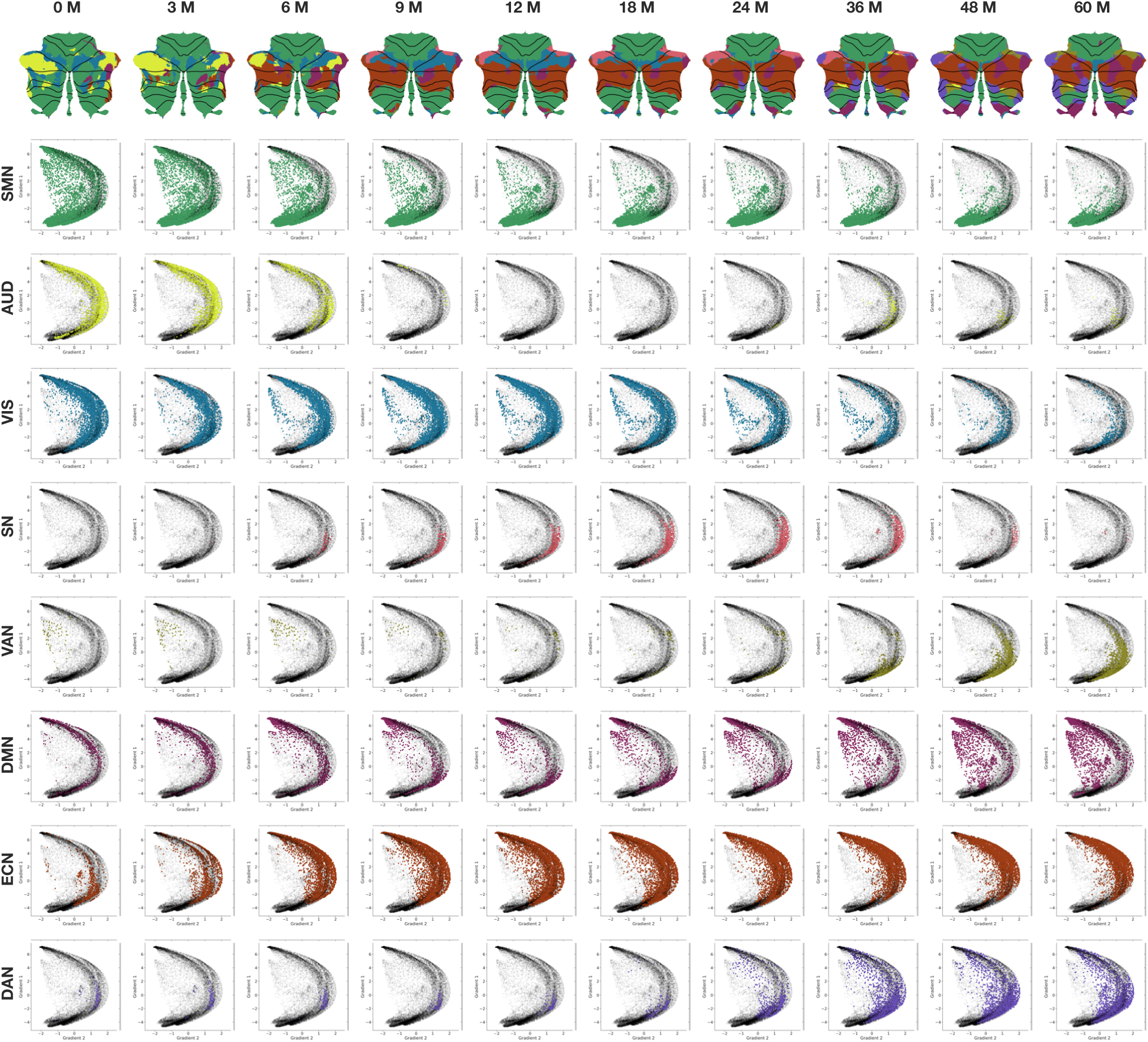
Functional gradients. Gradients of large-scale networks in children of both sexes over time.

**Fig. S10.**
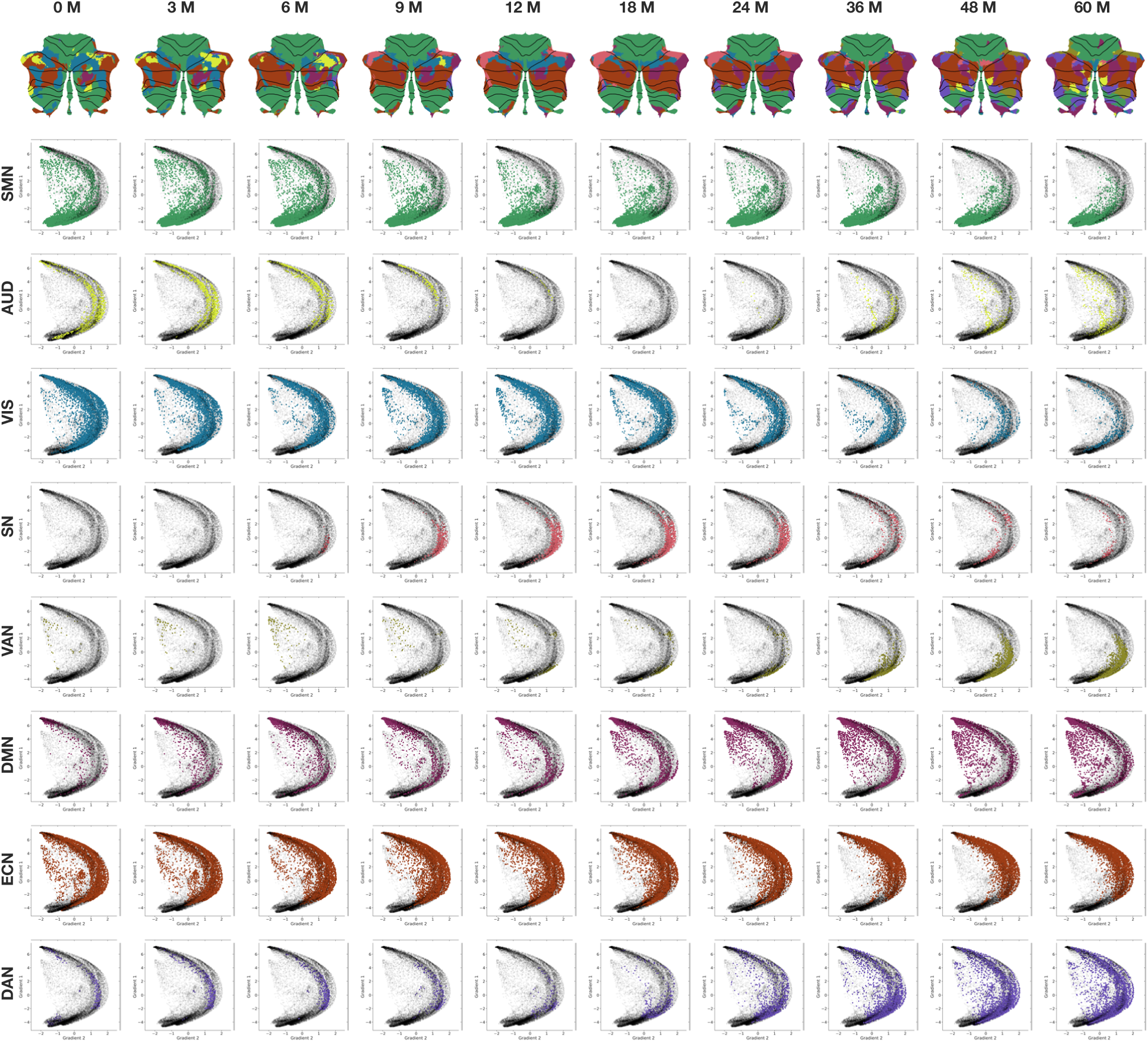
Functional gradients. Gradients of large-scale networks in female children over time.

**Fig. S11.**
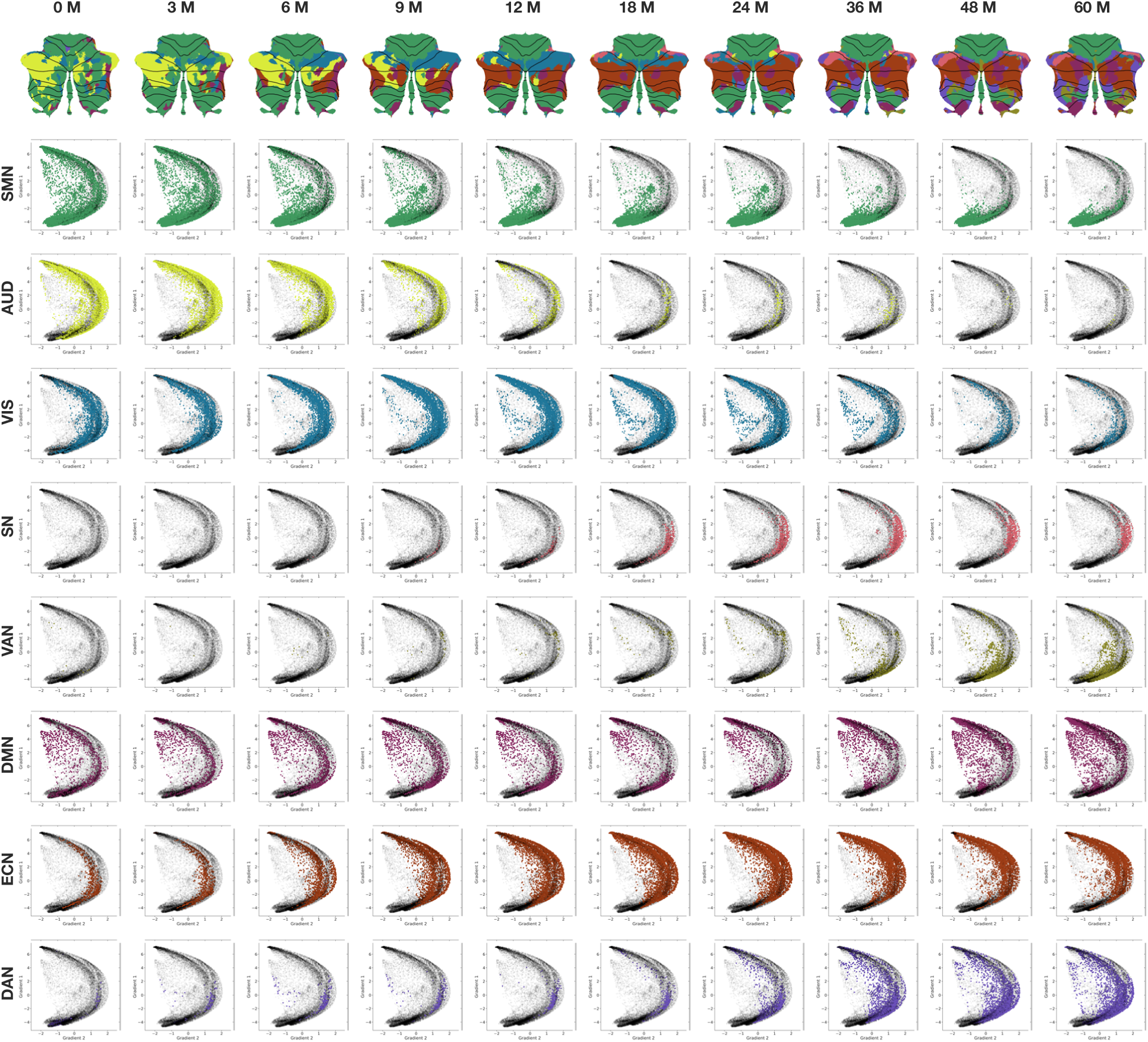
Functional gradients. Gradients of large-scales networks in male children over time.

**Tab. S2.**
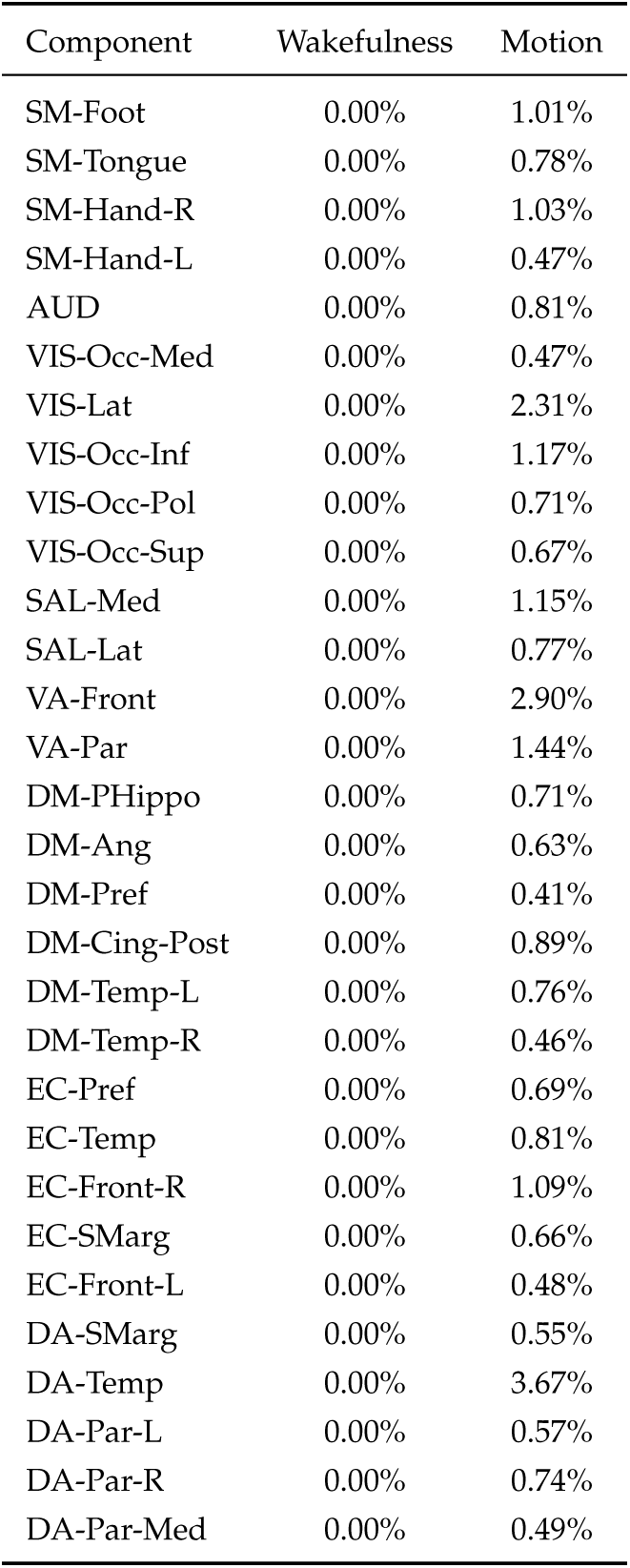
Effects of wakefulness and motion. Percentage of voxels with connectivity significantly affected (*p <* 0.01) by wakefulness and motion.

| Component | Wakefulness | Motion |
| --- | --- | --- |
| SM-Foot | 0.00% | 1.01% |
| SM-Tongue | 0.00% | 0.78% |
| SM-Hand-R | 0.00% | 1.03% |
| SM-Hand-L | 0.00% | 0.47% |
| AUD | 0.00% | 0.81% |
| VIS-Occ-Med | 0.00% | 0.47% |
| VIS-Lat | 0.00% | 2.31% |
| VIS-Occ-Inf | 0.00% | 1.17% |
| VIS-Occ-Pol | 0.00% | 0.71% |
| VIS-Occ-Sup | 0.00% | 0.67% |
| SAL-Med | 0.00% | 1.15% |
| SAL-Lat | 0.00% | 0.77% |
| VA-Front | 0.00% | 2.90% |
| VA-Par | 0.00% | 1.44% |
| DM-PHippo | 0.00% | 0.71% |
| DM-Ang | 0.00% | 0.63% |
| DM-Pref | 0.00% | 0.41% |
| DM-Cing-Post | 0.00% | 0.89% |
| DM-Temp-L | 0.00% | 0.76% |
| DM-Temp-R | 0.00% | 0.46% |
| EC-Pref | 0.00% | 0.69% |
| EC-Temp | 0.00% | 0.81% |
| EC-Front-R | 0.00% | 1.09% |
| EC-SMarg | 0.00% | 0.66% |
| EC-Front-L | 0.00% | 0.48% |
| DA-SMarg | 0.00% | 0.55% |
| DA-Temp | 0.00% | 3.67% |
| DA-Par-L | 0.00% | 0.57% |
| DA-Par-R | 0.00% | 0.74% |
| DA-Par-Med | 0.00% | 0.49% |

**Tab. S3.** Correlation with Mullen scores. Pearson correlations between the peak cerebellar connectivity with each cortical network and Mullen scores, adjusted for age using linear mixed models. Significant correlations (*p <* 0.05) are highlighted in bold. At coarse granularity, cerebellar connectivity with primary networks negatively correlates with fine motor scores, whereas connectivity with higher-order networks positively correlates with expressive language scores. At medium granularity, cerebellar connectivity with the SMN negatively correlates with fine motor scores, connectivity with the VAN negatively correlates with receptive language scores, and connectivity with the DAN positively correlates with both visual reception and expressive language scores.

| Network | Gross Motor | Fine Motor | Visual Reception | Expressive Language | Receptive Language |
| --- | --- | --- | --- | --- | --- |
| Primary | -0.056/0.158 | **-0.091/0.023** | -0.036/0.371 | 0.028/0.478 | -0.054/0.174 |
| Higher-Order | 0.018/0.652 | 0.004/0.912 | 0.018/0.650 | **0.105/0.008** | -0.055/0.169 |
| SMN | -0.030/0.457 | **-0.090/0.025** | -0.016/0.693 | -0.026/0.522 | -0.027/0.493 |
| AUD | -0.029/0.473 | -0.059/0.142 | 0.008/0.837 | 0.026/0.513 | -0.039/0.331 |
| VIS | -0.018/0.653 | -0.054/0.177 | -0.016/0.687 | 0.064/0.112 | -0.027/0.506 |
| SN | 0.015/0.711 | 0.000/0.995 | 0.022/0.586 | -0.044/0.274 | -0.017/0.679 |
| VAN | 0.016/0.686 | 0.036/0.368 | 0.051/0.207 | 0.034/0.399 | **-0.095/0.017** |
| DMN | 0.005/0.907 | -0.028/0.485 | 0.042/0.294 | 0.062/0.120 | -0.068/0.090 |
| ECN | 0.025/0.525 | -0.011/0.780 | -0.015/0.711 | 0.022/0.588 | -0.037/0.358 |
| DAN | 0.036/0.367 | -0.006/0.888 | **0.090/0.024** | **0.151/0.000** | -0.036/0.375 |

## References

1. Adamaszek, M. et al. Consensus paper: Cerebellum and emotion. The Cerebellum 16, 552–576 (2017). URL http://link.springer.com/10.1007/s12311-016-0815-8.

2. Hayter, A., Langdon, D. & Ramnani, N. Cerebellar contributions to working memory. NeuroImage 36, 943–954 (2007). URL https://linkinghub.elsevier.com/retrieve/pii/S1053811907001802.

3. Timmann, D. et al. The human cerebellum contributes to motor, emotional and cognitive associative learning. a review. Cortex 46, 845–857 (2010). URL https://linkinghub.elsevier.com/retrieve/pii/S0010945209002044.

4. Diedrichsen, J., King, M., Hernandez-Castillo, C., Sereno, M. & Ivry, R. B. Universal transform or multiple functionality? understanding the contribution of the human cerebellum across task domains. Neuron 102, 918–928 (2019). URL https://linkinghub.elsevier.com/retrieve/pii/S0896627319303782.

5. Barton, R. A. & Venditti, C. Rapid evolution of the cerebellum in humans and other great apes. Current Biology 24, 2440–2444 (2014). URL https://www.sciencedirect.com/science/article/pii/S0960982214010690.

6. Smaers, J. B. & Vanier, D. R. Brain size expansion in primates and humans is explained by a selective modular expansion of the cortico-cerebellar system. Cortex 118, 292–305 (2019). URL https://www.sciencedirect.com/science/article/pii/S0010945219301960.

7. Dellatolas, G. & Câmara-Costa, H. The role of cerebellum in the child neuropsychological functioning. In Handbook of Clinical Neurology, vol. 173, 265–304 (Elsevier, 2010). URL https://linkinghub.elsevier.com/retrieve/pii/B978044464150200023X.

8. Volpe, J. J. Cerebellum of the premature infant: Rapidly developing, vulnerable, clinically important. Journal of Child Neurology 24, 1085–1104 (2009). URL http://journals.sagepub.com/doi/10.1177/0883073809338067.

9. Limperopoulos, C., Chilingaryan, G., Guizard, N., Robertson, R. L. & Du Plessis, A. J. Cerebellar injury in the premature infant is associated with impaired growth of specific cerebral regions. Pediatric research 68, 145–150 (2010). URL https://www.nature.com/articles/pr2010148.

10. von Hofsten, C. & Rönnqvist, L. Preparation for grasping an object: a developmental study. Journal of experimental psychology: Human perception and performance 14, 610 (1988). URL 10.1037/0096-1523.14.4.610.

11. Butterworth, G., Verweij, E. & Hopkins, B. The development of prehension in infants: Halverson revisited. British Journal of Developmental Psychology 15, 223–236 (1997). URL 10.1111/j.2044-835X.1997.tb00736.x.

12. Karasik, L. B., Tamis-LeMonda, C. S. & Adolph, K. E. Transition from crawling to walking and infants’ actions with objects and people. Child Development 82, 1199–1209 (2011). URL https://srcd.onlinelibrary.wiley.com/doi/full/10.1111/j.1467-8624.2011.01595.x.

13. Adolph, K. E., Berger, S. E. & Leo, A. J. Developmental continuity? crawling, cruising, and walking. Developmental Science 14, 306–318 (2011). URL https://onlinelibrary.wiley.com/doi/full/10.1111/j.1467-7687.2010.00981.x.

14. Malina, R. M. Motor development during infancy and early childhood: Overview and suggested directions for research. International Journal of Sport and Health Science 2, 50–66 (2004). URL 10.5432/ijshs.2.50.

15. Gerber, R. J., Wilks, T. & Erdie-Lalena, C. Developmental milestones: motor development. Pediatrics in Review 31, 267–277 (2010). URL 10.1542/pir.31-7-267.

16. Tomasello, M. Language development. The Wiley-Blackwell handbook of childhood cognitive development 239–257 (2011). URL https://onlinelibrary.wiley.com/doi/pdf/10.1002/9781444325485#page=249.

17. Gilmore, J. H., Knickmeyer, R. C. & Gao, W. Imaging structural and functional brain development in early childhood. Nature Reviews Neuroscience 19, 123–137 (2018). URL https://www.nature.com/articles/nrn.2018.1.

18. Strüber, N. & Roth, G. Developmental Neurobiology, 117–141 (Springer Berlin Heidelberg, Berlin, Heidelberg, 2023). URL 10.1007/978-3-662-65774-4_5.

19. O’Reilly, J. X., Beckmann, C. F., Tomassini, V., Ramnani, N. & Johansen-Berg, H. Distinct and overlapping functional zones in the cerebellum defined by resting state functional connectivity. Cerebral Cortex 20, 953–965 (2010). URL 10.1093/cercor/bhp157.

20. Yeo, B. T. et al. The organization of the human cerebral cortex estimated by intrinsic functional connectivity. Journal of neurophysiology (2011). URL 10.1152/jn.00338.2011.

21. Sang, L. et al. Resting-state functional connectivity of the vermal and hemispheric subregions of the cerebellum with both the cerebral cortical networks and subcortical structures. NeuroImage 61, 1213–1225 (2012). URL https://linkinghub.elsevier.com/retrieve/pii/S1053811912003928.

22. Habas, C. et al. Distinct cerebellar contributions to intrinsic connectivity networks. Journal of Neuroscience 29, 8586–8594 (2009). URL https://www.jneurosci.org/content/29/26/8586.

23. Guell, X., Gabrieli, J. D. & Schmahmann, J. D. Triple representation of language, working memory, social and emotion processing in the cerebellum: convergent evidence from task and seed-based resting-state fMRI analyses in a single large cohort. NeuroImage 172 (2018). URL https://linkinghub.elsevier.com/retrieve/pii/S105381191830082X.

24. Guell, X., Schmahmann, J. D., Gabrieli, J. D. & Ghosh, S. S. Functional gradients of the cerebellum. eLife 7, e36652 (2018). URL https://elifesciences.org/articles/36652.

25. Guell, X. & Schmahmann, J. Cerebellar functional anatomy: a didactic summary based on human fMRI evidence. The Cerebellum 19, 1–5 (2020). URL https://link.springer.com/article/10.1007/s12311-019-01083-9.

26. Xue, A. et al. The detailed organization of the human cerebellum estimated by intrinsic functional connectivity within the individual. Journal of neurophysiology 125, 358–384 (2021). URL https://journals.physiology.org/doi/full/10.1152/jn.00561.2020.

27. Kipping, J. A., Tuan, T. A., Fortier, M. V. & Qiu, A. Asynchronous development of cerebellar, cerebello-cortical, and cortico-cortical functional networks in infancy, childhood, and adulthood. Cerebral Cortex 5170–5184 (2016). URL http://cercor.oxfordjournals.org/cgi/doi/10.1093/cercor/bhw298.

28. Diamond, A. Close interrelation of motor development and cognitive development and of the cerebellum and prefrontal cortex. Child Development 71, 44–56 (2000). URL https://srcd.onlinelibrary.wiley.com/doi/abs/10.1111/1467-8624.00117. https://srcd.onlinelibrary.wiley.com/doi/pdf/10.1111/1467-8624.00117.

29. Konrad, K. & Eickhoff, S. B. Is the ADHD brain wired differently? A review on structural and functional connectivity in attention deficit hyperactivity disorder. Human Brain Mapping 31, 904–916 (2010). URL https://onlinelibrary.wiley.com/doi/full/10.1002/hbm.21058.

30. Rubia, K. Cognitive neuroscience of attention deficit hyperactivity disorder (ADHD) and its clinical translation. Frontiers in Human Neuroscience 12, 100 (2018). URL http://journal.frontiersin.org/article/10.3389/fnhum.2018.00100/full.

31. Verly, M. et al. Altered functional connectivity of the language network in ASD: Role of classical language areas and cerebellum. NeuroImage: Clinical 4, 374–382 (2014). URL https://www.sciencedirect.com/science/article/pii/S2213158214000096.

32. Khan, A. J. et al. Cerebro-cerebellar resting-state functional connectivity in children and adolescents with autism spectrum disorder. Biological Psychiatry 78, 625–634 (2015). URL https://www.sciencedirect.com/science/article/pii/S0006322315002735.

33. Ramos, T. C., Balardin, J. B., Sato, J. R. & Fujita, A. Abnormal cortico-cerebellar functional connectivity in autism spectrum disorder. Frontiers in Systems Neuroscience 12 (2019). URL https://www.frontiersin.org/articles/10.3389/fnsys.2018.00074.

34. Adamaszek, M., Manto, M. & Schutter, D. J. L. G. (eds.) The Emotional Cerebellum, vol. 1378 of Advances in Experimental Medicine and Biology (Springer International Publishing, 2022). URL https://link.springer.com/10.1007/978-3-030-99550-8.

35. Howell, B. R. et al. The UNC/UMN baby connectome project (BCP): An overview of the study design and protocol development. NeuroImage 185, 891–905 (2019). URL https://linkinghub.elsevier.com/retrieve/pii/S1053811918302593.

36. Diedrichsen, J. A spatially unbiased atlas template of the human cerebellum. NeuroImage 33, 127–38 (2006). URL https://www.ncbi.nlm.nih.gov/pubmed/16904911.

37. Schmahmann, J. D. et al. Three-dimensional MRI atlas of the human cerebellum in proportional stereotaxic space. Neuroimage 10, 233–260 (1999). URL https://www.ncbi.nlm.nih.gov/pubmed/10458940.

38. Diedrichsen, J. & Zotow, E. Surface-based display of volume-averaged cerebellar imaging data. PLOS ONE 10, e0133402 (2015). URL https://dx.plos.org/10.1371/journal.pone.0133402.

39. van Es, D. M., van der Zwaag, W. & Knapen, T. Topographic maps of visual space in the human cerebellum. Current Biology 29, 1689–1694 (2019). URL 10.1016/j.cub.2019.04.012.

40. Buckner, R. L., Krienen, F. M., Castellanos, A., Diaz, J. C. & Yeo, B. T. T. The organization of the human cerebellum estimated by intrinsic functional connectivity. Journal of Neurophysiology 106, 2322–2345 (2011). URL https://journals.physiology.org/doi/full/10.1152/jn.00339.2011. Publisher: American Physiological Society.

41. King, M., Hernandez-Castillo, C. R., Poldrack, R. A., Ivry, R. B. & Diedrichsen, J. Functional boundaries in the human cerebellum revealed by a multi-domain task battery. Nature Neuroscience 22, 1371–1378 (2019). URL https://www.nature.com/articles/s41593-019-0436-x.

42. Guell, X., et al. LittleBrain: A gradient-based tool for the topographical interpretation of cerebellar neuroimaging findings. PLOS ONE 14, e0210028 (2019). URL https://journals.plos.org/plosone/article?id=10.1371/journal.pone.0210028.

43. Zhang, D. et al. Intrinsic functional relations between human cerebral cortex and thalamus. Journal of Neurophysiology 100, 1740–1748 (2008). URL https://www.physiology.org/doi/10.1152/jn.90463.2008.

44. Greene, D. J. et al. Developmental changes in the organization of functional connections between the basal ganglia and cerebral cortex. Journal of Neuroscience 34, 5842–5854 (2014). URL https://www.jneurosci.org/content/jneuro/34/17/5842.full.pdf.

45. Margulies, D. S. et al. Situating the default-mode network along a principal gradient of macroscale cortical organization. Proceedings of the National Academy of Sciences 113, 12574–12579 (2016). URL https://www.pnas.org/doi/full/10.1073/pnas.1608282113.

46. Wang, S. S.-H., Kloth, A. D. & Badura, A. The Cerebellum, Sensitive Periods, and Autism. Neuron 83, 518–532 (2014). URL https://www.cell.com/neuron/abstract/S0896-6273(14)00627-8.

47. Nettekoven, C., et al. A hierarchical atlas of the human cerebellum for functional precision mapping. Nature Communications 15 (2024). URL https://www.nature.com/articles/s41467-024-52371-w.

48. Yang, S. et al. The thalamic functional gradient and its relationship to structural basis and cognitive relevance. NeuroImage 218, 116960 (2020). URL https://linkinghub.elsevier.com/retrieve/pii/S1053811920304468.

49. Guell, X. Functional gradients of the cerebellum: A review of practical applications. The Cerebellum 21, 1061–1072 (2021). URL https://link.springer.com/10.1007/s12311-021-01342-8.

50. Menon, V. & Uddin, L. Q. Saliency, switching, attention and control: a network model of insula function. Brain Structure and Function 214, 655–667 (2010). URL 10.1007/s00429-010-0262-0.

51. Menon, V., Palaniyappan, L. & Supekar, K. Integrative brain network and salience models of psychopathology and cognitive dysfunction in schizophrenia. Biological Psychiatry 94, 108–120 (2023). URL https://linkinghub.elsevier.com/retrieve/pii/S0006322322016377.

52. Hu, D., Shen, H. & Zhou, Z. Functional asymmetry in the cerebellum: a brief review. The Cerebellum 7, 304–313 (2008). URL https://link.springer.com/article/10.1007/s12311-008-0031-2.

53. Fan, L. et al. Sexual dimorphism and asymmetry in human cerebellum: An MRI-based morphometric study. Brain Research 1353, 60–73 (2010). URL https://www.sciencedirect.com/science/article/pii/S0006899310015933.

54. Tiemeier, H. et al. Cerebellum development during childhood and adolescence: A longitudinal morphometric MRI study. NeuroImage 49, 63–70 (2010). URL https://linkinghub.elsevier.com/retrieve/pii/S1053811909008933.

55. Wang, Y. et al. Longitudinal development of the cerebellum in human infants during the first 800 days. Cell Reports 42, 112281 (2023). URL https://linkinghub.elsevier.com/retrieve/pii/S2211124723002929.

56. E, K., Chen, S. A., Ho, M. R. & Desmond, J. E. A meta-analysis of cerebellar contributions to higher cognition from PET and fMRI studies. Human Brain Mapping 35 (2012). URL https://www.ncbi.nlm.nih.gov/pmc/articles/PMC3866223/.

57. Marek, S. et al. Spatial and temporal organization of the individual human cerebellum. Neuron 100, 977–993.e7 (2018). URL https://linkinghub.elsevier.com/retrieve/pii/S0896627318308985.

58. Chechik, G., Meilijson, I. & Ruppin, E. Synaptic pruning in development: A computational account. Neural Computation 10, 1759–1777 (1998). URL https://direct.mit.edu/neco/article/10/7/1759-1777/6200.

59. Wiestler, T., McGonigle, D. J. & Diedrichsen, J. Integration of sensory and motor representations of single fingers in the human cerebellum. Journal of Neurophysiology 105, 3042–3053 (2011). URL https://journals.physiology.org/doi/full/10.1152/jn.00106.2011.

60. Paulin, M. G. The role of the cerebellum in motor control and perception. Brain Behavior and Evolution 41, 39–50 (1993). URL 10.1159/000113822.

61. Glickstein, M. What does the cerebellum really do? Current Biology 17, R824–R827 (2007). URL 10.1016/j.cub.2007.08.009.

62. Sasaki, K. Cerebro-cerebellar interactions and organization of a fast and stable hand movement: Cerebellar participation in voluntary movement and motor learning. In Cerebellar Functions, 70–85 (Springer, 1985).

63. Nawrot, M. & Rizzo, M. Motion perception deficits from midline cerebellar lesions in human. Vision Research 35, 723–731 (1995). URL https://linkinghub.elsevier.com/retrieve/pii/004269899400168L.

64. Wheelock, M. D. et al. Altered functional network connectivity relates to motor development in children born very preterm. NeuroImage 183, 574–583 (2018). URL https://www.sciencedirect.com/science/article/pii/S1053811918307493.

65. Ren, J. et al. Dissociable auditory cortico-cerebellar pathways in the human brain estimated by intrinsic functional connectivity. Cerebral Cortex 31, 2898–2912 (2021). URL https://academic.oup.com/cercor/article/31/6/2898/6120329.

66. Fransson, P., Åden, U., Blennow, M. & Lagercrantz, H. The functional architecture of the infant brain as revealed by resting-state fMRI. Cerebral Cortex 21, 145–154 (2010). URL 10.1093/cercor/bhq071. https://academic.oup.com/cercor/article-pdf/21/1/145/17304017/bhq071.pdf.

67. Pieterman, K. et al. Cerebello-cerebral connectivity in the developing brain. Brain Structure and Function 222, 1625–1634 (2017). URL https://link.springer.com/article/10.1007/s00429-016-1296-8.

68. Dijkshoorn, A. B. et al. Preterm infants with isolated cerebellar hemorrhage show bilateral cortical alterations at term equivalent age. Scientific Reports 10, 5283 (2020). URL https://www.nature.com/articles/s41598-020-62078-9.

69. Brossard-Racine, M., Du Plessis, A. J. & Limperopoulos, C. Developmental cerebellar cognitive affective syndrome in ex-preterm survivors following cerebellar injury. The Cerebellum 14, 151–164 (2015). URL https://link.springer.com/article/10.1007/s12311-014-0597-9.

70. Hortensius, L. M., et al. Neurodevelopmental consequences of preterm isolated cerebellar hemorrhage: A systematic review. Pediatrics 142 (2018). URL https://publications.aap.org/pediatrics/article/142/5/e20180609/38536/Neurodevelopmental-Consequences-of-Preterm.

71. Raz, G. & Saxe, R. Learning in infancy is active, endogenously motivated, and depends on the prefrontal cortices. Annual Review of Developmental Psychology 2, 247–268 (2020). URL 10.1146/annurev-devpsych-121318-084841.

72. Lyu, W., Wu, Y., Huynh, K. M., Ahmad, S. & Yap, P.-T. A multimodal submillimeter MRI atlas of the human cerebellum. Scientific Reports 14, 5622 (2024). URL https://www.nature.com/articles/s41598-024-55412-y.

73. Okayasu, M. et al. The Stroop effect involves an excitatory–inhibitory fronto-cerebellar loop. Nature Communications 14, 27 (2023). URL https://www.nature.com/articles/s41467-022-35397-w.

74. Sydnor, V. J. et al. Neurodevelopment of the association cortices: Patterns, mechanisms, and implications for psychopathology. Neuron 109, 2820–2846 (2021). URL https://linkinghub.elsevier.com/retrieve/pii/S0896627321004578.

75. Keller, A. S. et al. Personalized functional brain network topography is associated with individual differences in youth cognition. Nature Communications 14, 8411 (2023). URL https://www.nature.com/articles/s41467-023-44087-0.

76. Luo, A. C., et al. Functional connectivity development along the sensorimotor-association axis enhances the cortical hierarchy. Nature Communications 15 (2024). URL https://www.nature.com/articles/s41467-024-47748-w.

77. Wang, Y. et al. Spatio-molecular profiles shape the human cerebellar hierarchy along the sensorimotor-association axis. Cell Reports 43, 113770 (2024). URL https://linkinghub.elsevier.com/retrieve/pii/S2211124724000986.

78. Huntenburg, J. M., Bazin, P.-L. & Margulies, D. S. Large-scale gradients in human cortical organization. Trends in Cognitive Sciences 22, 21–31 (2018). URL https://www.cell.com/trends/cognitive-sciences/abstract/S1364-6613(17)30240-1.

79. Xia, Y. et al. Development of functional connectome gradients during childhood and adolescence. Science Bulletin 67, 1049–1061 (2022). URL https://www.sciencedirect.com/science/article/pii/S2095927322000020.

80. Taylor, H. P., et al. Functional hierarchy of the human neocortex from cradle to grave (2024). URL https://www.biorxiv.org/content/10.1101/2024.06.14.599109v1.

81. Scott, R. B. et al. Lateralized cognitive deficits in children following cerebellar lesions. Developmental Medicine & Child Neurology 43, 685–691 (2001). URL 10.1111/j.1469-8749.2001.tb00142.x.

82. Starowicz-Filip, A. et al. Cerebellar functional lateralization from the perspective of clinical neuropsychology. Frontiers in Psychology 12, 775308 (2021). URL https://www.frontiersin.org/journals/psychology/articles/10.3389/fpsyg.2021.775308/full.

83. Zahn-Waxler, C., Crick, N. R., Shirtcliff, E. A. & Woods, K. E. The origins and development of psychopathology in females and males. In Developmental Psychopathology, 76–138 (John Wiley & Sons, Ltd, 2015). URL https://onlinelibrary.wiley.com/doi/abs/10.1002/9780470939383.ch4.

84. Kucyi, A., Hove, M. J., Biederman, J., Van Dijk, K. R. & Valera, E. M. Disrupted functional connectivity of cerebellar default network areas in attention-deficit/hyperactivity disorder. Human Brain Mapping 36, 3373–3386 (2015). URL https://onlinelibrary.wiley.com/doi/full/10.1002/hbm.22850.

85. Özçalışkan, Ş. & Goldin-Meadow, S. Sex differences in language first appear in gesture. Developmental science 13, 752–760 (2010). URL https://onlinelibrary.wiley.com/doi/10.1111/j.1467-7687.2009.00933.x.

86. Tse, S. K., Chan, C., Li, H. & Kwong, S. M. Sex differences in syntactic development: Evidence from cantonese-speaking preschoolers in hong kong. International Journal of Behavioral Development 26, 509–517 (2002). URL https://journals.sagepub.com/doi/pdf/10.1080/01650250143000463.

87. Tomasi, D. & Volkow, N. D. Effects of family income on brain functional connectivity in US children: associations with cognition. Molecular Psychiatry 28, 4195–4202 (2023). URL https://www.nature.com/articles/s41380-023-02222-9.

88. Zhang, H., Shen, D. & Lin, W. Resting-state functional MRI studies on infant brains: A decade of gap-filling efforts. NeuroImage 185, 664–684 (2019). URL https://linkinghub.elsevier.com/retrieve/pii/S1053811918305962.

89. Jenkinson, M., Bannister, P., Brady, M. & Smith, S. Improved optimization for the robust and accurate linear registration and motion correction of brain images. NeuroImage 17, 825–841 (2002). URL https://www.sciencedirect.com/science/article/pii/S1053811902911328.

90. Andersson, J. L., Skare, S. & Ashburner, J. How to correct susceptibility distortions in spin-echo echo-planar images: application to diffusion tensor imaging. NeuroImage 20, 870–888 (2003). URL https://www.sciencedirect.com/science/article/pii/S1053811903003367.

91. Smith, S. M., et al. Advances in functional and structural MR image analysis and implementation as FSL. NeuroImage 23, S208–S219 (2004). URL https://www.sciencedirect.com/science/article/pii/S1053811904003933.

92. Greve, D. N. & Fischl, B. Accurate and robust brain image alignment using boundary-based registration. NeuroImage 48, 63–72 (2009). URL 10.1016/j.neuroimage.2009.06.060.

93. Glasser, M. F. et al. A multi-modal parcellation of human cerebral cortex. Nature 536, 171–178 (2016). URL https://www.nature.com/articles/nature18933.

94. Pruim, R. H. et al. Ica-aroma: A robust ica-based strategy for removing motion artifacts from fmri data. NeuroImage 112, 267–277 (2015). URL 10.1016/j.neuroimage.2015.02.064.

95. Parkes, L., Fulcher, B., Yücel, M. & Fornito, A. An evaluation of the efficacy, reliability, and sensitivity of motion correction strategies for resting-state functional MRI. NeuroImage 171, 415–436 (2018). URL 10.1016/j.neuroimage.2017.12.073.

96. Fonov, V. S., Evans, A. C., McKinstry, R. C., Almli, C. R. & Collins, D. Unbiased nonlinear average age-appropriate brain templates from birth to adulthood. NeuroImage 47, S102 (2009). URL 10.1016/S1053-8119(09)70884-5.

97. Fonov, V. et al. Unbiased average age-appropriate atlases for pediatric studies. Neuroimage 54, 313–327 (2011). URL 10.1016/j.neuroimage.2010.07.033.

98. Isensee, F., Jaeger, P. F., Kohl, S. A., Petersen, J. & Maier-Hein, K. H. nnu-net: a self-configuring method for deep learning-based biomedical image segmentation. Nature methods 18, 203–211 (2021). URL 10.1038/s41592-020-01008-z.

99. Glasser, M. F. et al. The minimal preprocessing pipelines for the human connectome project. Neuroimage 80, 105–124 (2013). URL 10.1016/j.neuroimage.2013.04.127.

100. Power, J. D., Barnes, K. A., Snyder, A. Z., Schlaggar, B. L. & Petersen, S. E. Spurious but systematic correlations in functional connectivity MRI networks arise from subject motion. NeuroImage 59, 2142–2154 (2012). URL 10.1016/j.neuroimage.2011.10.018.

101. Smith, S. M. et al. Correspondence of the brain’s functional architecture during activation and rest. Proceedings of the National Academy of Sciences 106, 13040–13045 (2009). URL http://www.pnas.org/cgi/doi/10.1073/pnas.0905267106.

102. Beckmann, C., Mackay, C., Filippini, N. & Smith, S. Group comparison of resting-state FMRI data using multi-subject ICA and dual regression. NeuroImage 47, S148 (2009). URL https://linkinghub.elsevier.com/retrieve/pii/S1053811909715113.

103. Greicius, M. D., Krasnow, B., Reiss, A. L. & Menon, V. Functional connectivity in the resting brain: a network analysis of the default mode hypothesis. Proceedings of the National Academy of Sciences 100, 253–258 (2003). URL https://www.pnas.org/doi/abs/10.1073/pnas.0135058100.

104. Yuen, N. H., Osachoff, N. & Chen, J. J. Intrinsic frequencies of the resting-state fMRI signal: the frequency dependence of functional connectivity and the effect of mode mixing. Frontiers in Neuroscience 13, 463704 (2019). URL https://www.frontiersin.org/journals/neuroscience/articles/10.3389/fnins.2019.00900/full.

105. Bro, R., Acar, E. & Kolda, T. G. Resolving the sign ambiguity in the singular value decomposition. Journal of Chemometrics: A Journal of the Chemometrics Society 22, 135–140 (2008). URL https://analyticalsciencejournals.onlinelibrary.wiley.com/doi/epdf/10.1002/cem.1122.

106. Wood, S. & Scheipl, F. gamm4: Generalized additive mixed models using ‘mgcv’ and ‘lme4’ (2017). URL https://CRAN.R-project.org/package=gamm4. R package version 0.2-6.

107. Spisák, T. et al. Probabilistic TFCE: A generalized combination of cluster size and voxel intensity to increase statistical power. NeuroImage 185, 12–26 (2019). URL https://www.sciencedirect.com/science/article/pii/S1053811918319505.

108. Mullen, E. M. et al. Mullen scales of early learning (AGS Circle Pines, MN, 1995).

109. Wickham, H. ggplot2: Elegant Graphics for Data Analysis (Springer-Verlag New York, 2016). URL https://ggplot2.tidyverse.org.

